# EGFR meets hATG8s – Biophysical and structural insights supporting a unique role of GABARAP during receptor trafficking

**DOI:** 10.1101/2024.07.10.602929

**Authors:** Alina Üffing, Oliver H. Weiergräber, Melanie Schwarten, Silke Hoffmann, Dieter Willbold

## Abstract

The human ATG8 family member GABARAP is involved in numerous autophagy-related and - unrelated processes. We recently observed that specifically the deficiency of GABARAP enhances EGFR degradation upon ligand stimulation. Here, we report on two putative LC3-interacting regions (LIRs) within the EGFR, the first of which (LIR1) is selected as GABARAP binding site *in-silico*. Indeed, *in-vitro* interaction studies reveal preferential binding of LIR1 to GABARAP and GABARAPL1. Our X-ray data demonstrate interaction of core LIR1 residues FLPV with both hydrophobic pockets of GABARAP suggesting a canonical binding. Although LIR1 occupies the LIR docking site, GABARAP Y49 and L50 appear dispensable this case. Our data support the hypothesis that GABARAP affects the fate of EGFR at least in part through direct binding.

## Introduction

The human autophagy related protein 8 (hATG8) family consists of seven members, which can be divided into the GABARAP (GABARAP, GABARAPL1 and GABARAPL2) and LC3 (LC3A, LC3B, LC3B2 and LC3C) subfamilies. They are ubiquitin-like modifiers which can be conjugated to membranes in a E1-E2-E3 like enzyme cascade [1–3]. In the past, hATG8s have been most extensively studied for their role in the conserved catabolic process of macroautophagy, hereafter referred to as autophagy [4,5]. While they are involved during the different steps of autophagy, from phagophore formation and extension to cargo sequestration and fusion with the lysosome [6–9], it still remains to be elucidated whether and for which steps they are essential and where alternative mechanisms take place [10]. Additionally, conjugation of hATG8 to single endolysosomal membranes (CASM) has been reported as a key function in a variety of processes related to inflammation, cancer and neurodegeneration [11–14]. Differential binding of ATG16L1, a component of the E3-ligase complex for membrane conjugation of hATG8s, appears to be decisive for whether conjugation occurs on single or double membranes [15]. Interestingly, CASM has been shown to be important for receptor recycling of TREM2, CD36, and TLR4 in the context of cellular uptake of amyloid β [16] as well as secretion of the transferrin receptor in a process called LC3-dependent extracellular vesicle loading and secretion (LDELS) [17]. The trafficking of the γ-aminobutyric acid type A (GABA_A_) receptor, the eponymous GABARAP binding partner, to the plasma membrane is another example of a non-conventional role of a hATG8 [18,19]. In addition to membrane conjugation, the versatile functions of hATG8 proteins are facilitated through interactions with a conserved LC3-interacting region (LIR) [20], with its core consisting of four amino acids (Θ_0_-X_1_-X_2_-Γ_3_) with Θ being an aromatic residue (W/F/Y) and Γ an aliphatic residue (L/I/V). Both the residues of the core LIR as well as surrounding residues regulate selective binding to the GABARAP or LC3 subfamily proteins, with Unc-51-like kinase 1 (ULK1) being an example of a LIR-containing protein with high preference for GABARAP and GABARAPL1 (30-fold affinity compared to the LC3 subfamily proteins) [20,21]. Moreover, binding to hATG8s can be altered towards higher or lower affinities by phosphorylation [22]. Phosphorylation of the Golgi protein SCOC increases its affinity for the LC3 subfamily [23], and OPTN phosphorylation and subsequent increase in affinity has been shown to be important for autophagic clearance of *Salmonella* [24]. In contrast, tyrosine phosphorylation of the Θ_0_ residue of the mitophagy receptor FUNDC1 has been reported to weaken its interaction with LC3 [25]. Recently, we observed enhanced degradation of the epidermal growth factor (EGF)-receptor (EGFR) in response to EGF stimulation in cells deficient in GABARAP but not its paralogs, which was accompanied by decreased MAPK/ERK signaling as well as altered target gene expression [26]. Consistently, continuous live-cell imaging revealed lower EGF-647 levels in Huh-7.5 GABARAP single knockout cells compared to Huh-7.5 wildtype cells over time (Fig. S1A). Owing to two putative LIRs present in the regulatory C-terminal domain of the EGFR, we sought to investigate this phenotype from a biophysical and structural perspective, analyzing the specific binding mode shaping this putative interaction.

## Material and Methods

### DNA constructs

Genes encoding proteins for purification were expressed from pGEX-4-T2 vectors previously described [27,28], which can be found at Addgene for GABARAP (#73948), GABARAPL1 (#73945), GABARAPL2 (#73518) and LC3A (#73946). pGEX-4-T2-LC3B was cloned by restriction-ligation using BamHI and NotI. Point mutations substituting Y49 and L50 to alanines were introduced by site-directed mutagenesis for GABARAP and GABARAPL1. All constructs encoded full-length (unprocessed) versions of human ATG8 proteins. EGFR LIR1_1076–1099_ fused to GABARAP by a glycine-serine linker (hereafter only EGFR LIR1-GABARAP) was ordered as a codon-optimized, synthetic construct in pGEX-4T-2 from GeneArt (Thermo Fisher Scientific). Please note that our EGFR numbering refers to the full-length receptor including the signal peptide (first 24 aa).

### Protein expression and purification

GABARAP, GABARAP_Y49A/L50A_, GABARAPL1, GABARAPL1_Y49A/L59A_, GABARAPL2, LC3A and LC3B used for biolayer interferometry (BLI) experiments were expressed and purified from *E. coli* BL21(DE3) transformed with respective pGEX-4-T2 plasmids as previously described [29]. In short, glutathione S-transferase (GST) fusion proteins were first purified from soluble extracts by affinity chromatography using Glutathione Sepharose 4B (GE Healthcare). After cleavage of the GST tag with thrombin, further purification was carried out by size exclusion chromatography using either a Hiload 26/60 Superdex 75 or a Hiload Superdex 16/600 75 preparatory grade column, which were equilibrated with 25 mM Tris, 150 mM NaCl (pH 7.5) and 0.5 mM tris(2-carboxyethyl)phosphine (TCEP), if proteins contained Cysteine, and eluted in the same buffer. GST-EGFR-LIR1-GABARAP protein for X-ray crystallography was purified analogously from *E. coli* BL21(DE3) transformed with pGEX-4-T2-EGFR LIR1-GABARAP grown in LB medium. Gene expression was induced with 1 mM isopropyl β-d-1-thiogalactopyranoside (IPTG) at an OD_600nm_ of 0.7 and for 22 hat 25 °C. Afterwards, cells were harvested by centrifugation at 3000 × g for 30 min at 4 °C and washed once with PBS. The GST fusion protein was purified from the soluble extract by affinity chromatography using Glutathione Sepharose 4B (GE Healthcare). The GST-tag was cleaved off using thrombin, yielding a 145 aa EGFR LIR1-GABARAP fusion, which was further purified by size exclusion chromatography using a Hiload 26/60 Superdex 75 preparatory grade column equilibrated with 10 mM Tris-HCl, 150 mM sodium chloride, pH 7 and eluted in the same buffer. ^15^N-GABARAP for Heteronuclear Single Quantum Coherence (HSQC)-titration experiments was purified from *E. coli* BL21(DE3) transformed with pGEX-4-T2-GABARAP grown in M9 minimal medium with 1g/L ^15^NH_4_Cl. Gene expression was induced with 1 mM IPTG at an OD_600nm_ of 0.7 and for 20 h at 25 °C. Cells were harvested and ^15^N-GST-GABARAP was purified by affinity chromatography as described above. After thrombin cleavage, ^15^N-GABARAP was further purified by size exclusion chromatography using a Hiload 26/60 Superdex 75 preparatory grade column equilibrated with 25 mM NaH_2_PO_4_/Na_2_HPO_4_, 100 mM KCl, 100 mM NaCl, 50 µM ethylenediaminetetraacetic acid (EDTA), pH 6.9 and eluted in the same buffer.

### Biolayer interferometry

BLI experiments were performed on an Octet Red 96 (FORTÉBIO) as previously described [26]. In short, peptides with N-terminal biotinylation were ordered from CASLO (≥95% purity, SI Table 1) and immobilized on High Precision Streptavidin (SAX) biosensors (FORTÉBIO/Sartorius). Purified hATG8 proteins were used as analyte in increasing concentrations (Supplementary data). For dissociation constant (K_D_) calculation, respective reference sensor response levels were subtracted and baselines aligned followed by steady-state evaluation by plotting the respective response levels against the applied protein concentration. Curves were fitted by non-linear regression according to the One-site binding model using GraphPad Prism version 9 for Windows (GraphPad Software, Boston, MA, USA). Only fitted K_D_ values corresponding to a saturation level of at least 0.25 nm and R-square above 0.985 are shown. Quality of each hATG8 as an analyte was assessed by also determining its binding to the PCM1 LIR peptide, for which published reference data are available w.r.t. all hATG8s [20]. Owing to the many different ligand-analyte combinations, we report the results of individual measurements, unless stated otherwise (refer to SI data related to Fig.1B, 1C, 4B, 4C, S1C).

**Figure 1.**
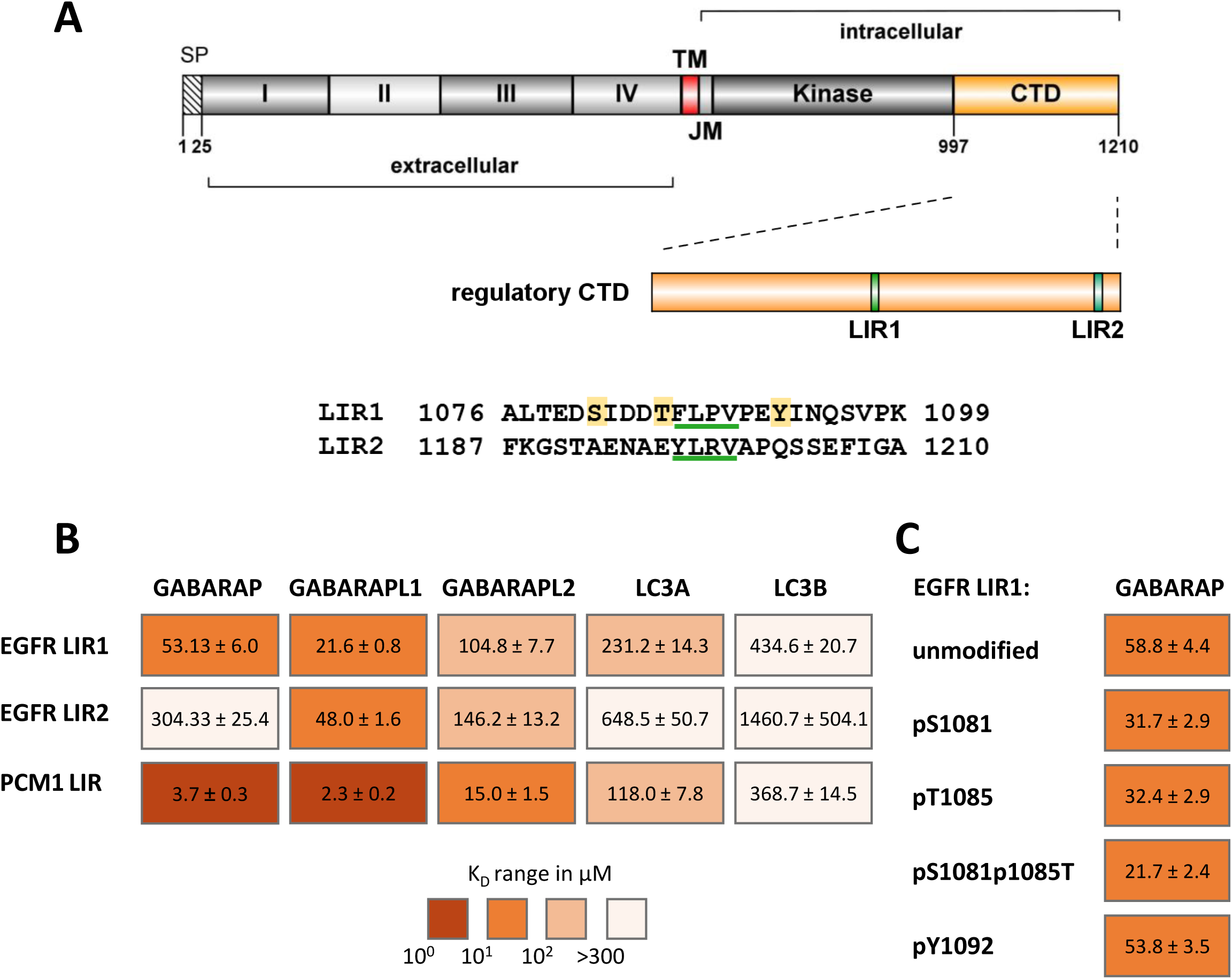
EGFR LIR1 preferentially interacts with GABARAP/L1 with only moderate influence of phosphorylation (**A**) Schematic representation of the EGFR domain structure with LIR1 and LIR2 indicated in the C-terminal domain as well as the respective amino acid sequences. Core LIRs are underlined in green. For LIR1, phosphorylation sites investigated in (C) are shown in yellow. (**B**) K_D_ values [µM] of human ATG8 paralogs GABARAP, GABARAPL1, GABARAPL2, LC3A and LC3B with EGFR peptides. (**C**) K_D_ values [µM] of phosphorylated (p) EGFR LIR1 variants with GABARAP. Color scaled according to affinity as determined by biolayer interferometry. K_D_ values are shown with standard error calculated from non-linear regression. Abbreviations: SP (signal peptide), TM (transmembrane), JM (juxtamembrane). EGFR was illustrated using IBS 2.0 [35].

### X-ray crystallography and data processing

EGFR LIR1-GABARAP in 10 mM Tris-HCl and 150 mM NaCl, pH 7, was concentrated to approximately 17 mg/ml using Vivaspin 20/2 centrifugal filter units (3 kDa cutoff). Crystallization was performed by the sitting-drop vapor diffusion method, using a Freedom Evo robotic device (Tecan) with commercially available screening sets and combining 0.5 μl of protein solution and 0.5 μl of reservoir solution for each drop. Several conditions yielded crystals. The crystal which was used for structure determination developed in 0.17 M ammonium acetate, 0.085 M sodium citrate, pH 5.6, 25.5% (w/v) PEG 4000 and 15% (v/v) glycerol. X-ray diffraction data was collected at 100 K on beamline ID30A-3/MASSIF-3 of the European Synchrotron Radiation Facility (ESRF; Grenoble, France; doi: 10.15151/ESRF-DC-1524662410). XDS and XSCALE [30] were used for data processing and reflections to a d_min_ of 2.05 Å were included in the final dataset. The structure of GABARAP was determined by molecular replacement with MOLREP [31] using the structure of GABARAP from its K1 peptide complex (PDB ID: 3D32) as template. Coordinates of the EGFR LIR1 peptide were generated using COOT [32], and the model was improved by reciprocal-space refinement with phenix.refine [33] alternating with interactive rebuilding in COOT. According to validation using the wwPDB validation pipeline, the model features good geometry with no outliers in the Ramachandran plot and less than 1% of unusual side chain rotamers. Coordinates and structure factor amplitudes were deposited in the PDB (www.ebi.ac.uk/pdbe) with accession number 8S1M (doi: 10.2210/pdb8S1M/pdb). For statistics of data collection and refinement, refer to Supplementary Table 2. Figures were created using the PyMOL Molecular Graphics System, Version 3.0, Schrödinger, LLC.

### NMR titrations and data analysis

Titrations of GABARAP with EGFR LIR1 and PCM1 LIR peptides spanning residues 1076–1099 and 1954–1968, respectively, were monitored by recording 2D [^1^H,^15^N] HSQC spectra at a temperature of 25.0°C on a Bruker 900 MHz Avance Neo spectrometer equipped with a triple resonance ^1^H, ^13^C, ^15^N TCI cryoprobe. ^15^N-GABARAP was concentrated to 200 µM in 25 mM NaH_2_PO_4_/Na_2_HPO_4_, 100 mM KCl, 100 mM NaCl, 50 µM EDTA, and 5% (v/v) D_2_O. 2D [^1^H,^15^N] HSQC spectra were recorded after stepwise addition of peptide up to a two-fold molar excess. For chemical shift perturbation (CSP) analysis, ^1^H chemical shift changes, Δδ(^1^H), and ^15^N chemical shift changes, Δδ(^15^N), in units of ppm were combined according to the following equation:

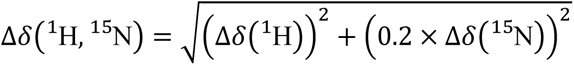

For mapping on GABARAP (PDB ID: 1KOT), residues were colored according to Δδ(^1^H, ^15^N) values at twofold molar excess using the PyMOL Molecular Graphics System, Version 3.0 Schrödinger, LLC.

## Results

### EGFR LIR1 preferentially interacts with GABARAP and GABARAPL1 with minor impact of phosphorylation

The previously reported enhanced EGFR degradation after EGF stimulation, specifically in GABARAP single knockout cells included a first hypothesis regarding a direct interaction between GABARAP and EGFR, including the observation of a putative extended LIR motif (positions 1086–1089) in the regulatory carboxyterminal domain (CTD) of EGFR [26]. Interestingly, the regulatory C-terminal domain (CTD) of EGFR, exogenously expressed as a fusion protein with green fluorescent protein (EGFR CTD-GFP) in HEK293 cells, can be enriched from whole cell lysates by purified GST-GABARAP (Fig. S1B). Moreover, by the iLIR prediction tool [34] another putative LIR motif located adjacent to the very end of the CTD is predicted (Fig. 1A). While the first putative core LIR comprises the amino acids FLPV, the second (position 1197 to 1200) consists of YLRV. Considering the EGFR sequences of different mammals, segments corresponding to LIR1 and LIR2 appear to range among the more conserved regions within the CTD, with frequently identical core sequences (Fig. S2); this suggests that an activity of these regions is preserved during evolution, a fact that often applies to functional LIRs [36]. We next tested whether these two LIRs are also selected as GABARAP binding sites within EGFR CTD-derived sequences by the artificial intelligence-based structure prediction tool AlphaFold 2 (AF2)-multimer [37,38], which can predict binding sites of ATG8s with high accuracy especially in intrinsically disordered protein regions [36]. While we found FLPV-related binding in many of the resulting models, not a single model involved YLRV (Fig. S3A and B). In the following, we used synthetic peptides, representing residues 1076–1099 (LIR1) and 1190–1210 (LIR2) of EGFR for *in-vitro* binding experiments with GABARAP applying biolayer interferometry (BLI, for peptide sequences refer to SI Table 1). To investigate whether the observed GABARAP-specific phenotype previously observed in HEK293 and Huh-7.5 cells might be mirrored by a direct and paralog-specific interaction, we included all paralogs with notable mRNA expression levels (cutoff: nTPM >10, for values and details refer to SI Table 3) in at least one of the two cell lines, these being GABARAP, GABARAPL1, GABARAPL2, LC3A and LC3B, in our analysis A peptide comprising the LIR motif (residues 1954-1968) of the well-described GABARAP interactor pericentriolar material 1 protein (PCM1) was included as control, with our K_D_ values largely matching those published [20]. When comparing LIR1 and LIR2 binding, a clear difference was found especially for GABARAP, with an about 6-fold higher affinity for LIR1 (K_D_ of 54.7 +/− 3.6 µM (mean/SD from three independent experiments)) over LIR2, while for the other investigated paralogs affinity differences were more modest (about 1.4 to 3.4-fold higher for LIR1), with LIR2 binding usually still appearing weaker. Importantly, LIR1 showed a particular specificity for GABARAP and its closest relative, GABARAPL1, with approximately 2-fold (5-fold), 4-fold (11-fold) and 8-fold (20-fold) stronger binding of GABARAP (GABARAPL1) to LIR1 relative to the respective affinities obtained for the paralogs GABARAPL2, LC3A and LC3B. For both LIR peptides, the lowest affinities were measured for LC3B with 3-digit µM to mM K_D_ values (Fig. 1B). Interestingly, both LIR peptides contain phosphorylation sites, of which Y1092 and Y1197 have been extensively described for their role in EGFR signaling [39–44] while the roles of S1081 and T1085 remain more elusive [45–47]. Phosphorylation of either S1081 or T1085, both located upstream of the core LIR1, slightly enhanced GABARAP’s affinity (∼1.8-fold), while phosphorylation of both sites simultaneously enhanced its binding to above two-fold change (2.7-fold, Fig. 1C). Our *in-vitro* interaction studies revealed virtually unchanged binding of EGFR LIR1 to GABARAP when phosphorylated at residue Y1092 located downstream of the core LIR1, which also applies for the paralogs (Fig. S4, top). Curiously, phosphorylation of Θ_0_ residue Y1197 within the core (YLRV) of LIR2 did not show the expected detrimental effect on the interaction of LIR2 with GABARAP or with its paralogs (Fig. S4, bottom).

As GABARAPL1 exhibits by far lower expression levels than GABARAP (www.proteinatlas.org, v23; [48]) within the cell lines analyzed in Dobner (2020), our structural investigations focused on the interaction of EGFR LIR1 with GABARAP.

### X-ray complex structure reveals canonical binding of EGFR LIR1 to GABARAP

To further investigate the interaction mode between the putative LIR1 of the EGFR and GABARAP, a chimeric protein consisting of EGFR residues 1076–1099 fused to GABARAP (EGFR LIR1-GABARAP) was expressed, purified and used for crystallization. We were able to obtain crystals and determine the structure, resolving GABARAP together with residues 1082–1099 of the EGFR LIR1 peptide (Fig. 2A, SI Table 1), which were found to interact with the GABARAP moiety of a symmetry-equivalent molecule. While the two N-terminal helices and the ubiquitin-like core of GABARAP as well as the core LIR1 of EGFR (FLPV), which contacts the hydrophobic pockets of GABARAP, featured well defined electron density, EGFR residues surrounding the core LIR were less clearly defined. Together with elevated B factors obtained during refinement, this points towards enhanced dynamics of both segments flanking the core LIR (Fig. 2B). Consequently, the following structural analysis of the inter-molecular interface between GABARAP and EGFR LIR1 focusses on the core LIR. Residue F1086 of the EGFR, representing position Θ_0_ of the core LIR motif, is inserted into hydrophobic pocket 1 (HP1), supported by hydrophobic interactions with residues I21, P30, L50, K48 and F104 of GABARAP. The side chain of core LIR residue X_1_ (L1087) is within van der Waals distance of Y49 and K46, possibly involved in hydrophobic interactions, as well as R67. Additionally, the amide nitrogen and carbonyl oxygen of EGFR L1087 are oriented towards strand β2 of GABARAP and placed within hydrogen bonding distance of its K48 carbonyl oxygen and its L50 amide, respectively. In contrast, EGFR residue P1088 (X_2_) does not appear to be involved in the interaction with GABARAP. The second hydrophobic pocket of GABARAP is occupied by V1089 of EGFR, representing Γ_3_ of the core LIR motif, which is involved in hydrophobic interactions with V51 and P52 (Fig. 2C). Superposition with the complex structure of the ULK1 LIR with GABARAP [20] reveals a similar binding interface of ULK1 residues 355 to 366 and EGFR residues 1084 to 1095 with GABARAP (Fig. 2D). While Θ_0_ and Γ_3_ as well as the X_4_ position are represented by the same amino acids (F, V, and P, respectively), X_1_ and X_2_ differ between the two peptides. The ULK1 LIR possesses a valine at the X_1_ position, which likely allows for similar hydrophobic interactions with Y49 as L1087 in the EGFR LIR1. However, while P1088 in X_2_ shows no direct contact with GABARAP, the corresponding M359 in the ULK1 LIR is positioned in close proximity to GABARAP residues Y25 and L50 and thus interacts with HP1. This could partially explain the much stronger (∼1000-fold, [20]) binding affinity of the ULK1 peptide, despite the similarity of the LIRs. We also superimposed our complex structure with the *in-silico* structural models and found a high degree of agreement between the relevant binding interfaces regardless of whether the complete cytoplasmic domain of the EGFR or shorter fragments thereof served as input (Fig. S3B), confirming the previously described accurate predictability of ATG8-ligand complexes by AF2 [36].

**Figure 2.**
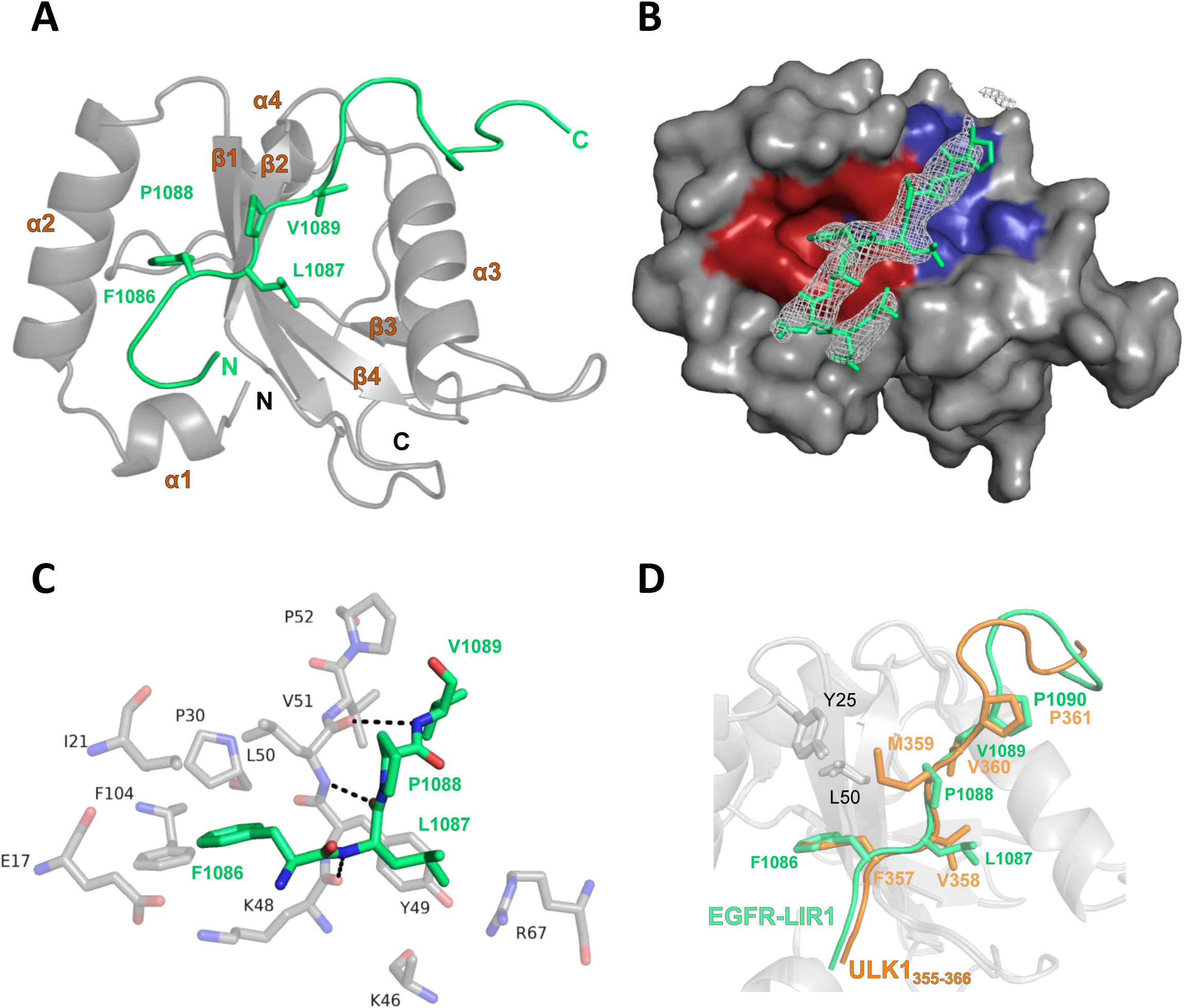
X-ray structure of EGFR LIR1-GABARAP. (**A**) Overall structure of the EGFR LIR1 in complex with GABARAP presented as cartoon. GABARAP is shown in gray and the EGFR peptide in green. Coordinates and structure factors have been deposited in the Protein Data Bank with accession number 8S1M. (**B**) Surface representation of GABARAP with HP1 (red) and HP2 (blue) indicated. The 2mF_o_−DF_c_ map covering the EGFR peptide is colored white and contoured at 0.8 sigma and the EGFR peptide model is shown in green. (**C**) Core LIR residues (FLPV, in green) together with respective GABARAP residues at a distance lower than 4 Å. Hydrogen bonds are displayed as dashed lines. (**D**) Comparison of EGFR LIR1 and ULK1 extended core LIR (PDB: 6HYO, [20]) with GABARAP.

### EGFR and PCM1 LIR peptides impact the same region of GABARAP in solution

In order to gain more precise information on the contribution of individual GABARAP residues, we next mapped the GABARAP binding interface of a free EGFR LIR peptide using NMR chemical shift perturbation (CSP) data, and in addition compared it with that of the PCM1 LIR. As seen in the 2D ^1^H-^15^N HSQC spectrum (Fig. 3A), GABARAP without ligand exhibited the known resonances for natively folded GABARAP. Upon stepwise addition of the EGFR LIR1 ligand, several peaks displayed changes in chemical shift or line broadening beyond detection. At twofold molar excess of EGFR LIR1 peptide, residues surrounding the hydrophobic pockets, HP1 and HP2 of GABARAP, specifically I21, R22, V29, V31, K46, Y49, L50, S53, F60 and L63 showed chemical shift perturbations above 0.1 ppm. Additionally, the peaks for Y25, K48 and V51 had disappeared, indicating intermediate exchange and thus a defining character of these residues for EGFR LIR1 binding (Fig. 3B, Fig. S5). In comparison, more residues were affected with the PCM1 LIR peptide as ligand at two-fold excess, with more residue peaks showing line broadening beyond detection (Fig. 3C, Fig. S5). This included the LDS residues Y49 and L50 of GABARAP, which have been previously described to be crucial for many LIR-LDS interactions [49]. Overall, the mapping of the CSP data on the GABARAP surface shows that binding of both ligands, EGFR LIR1 and PCM1 LIR, affect the same region, namely the LDS of GABARAP in a canonical manner, however, Y49 and L50 appeared to be less defining for binding of EGFR LIR1.

**Figure 3.**
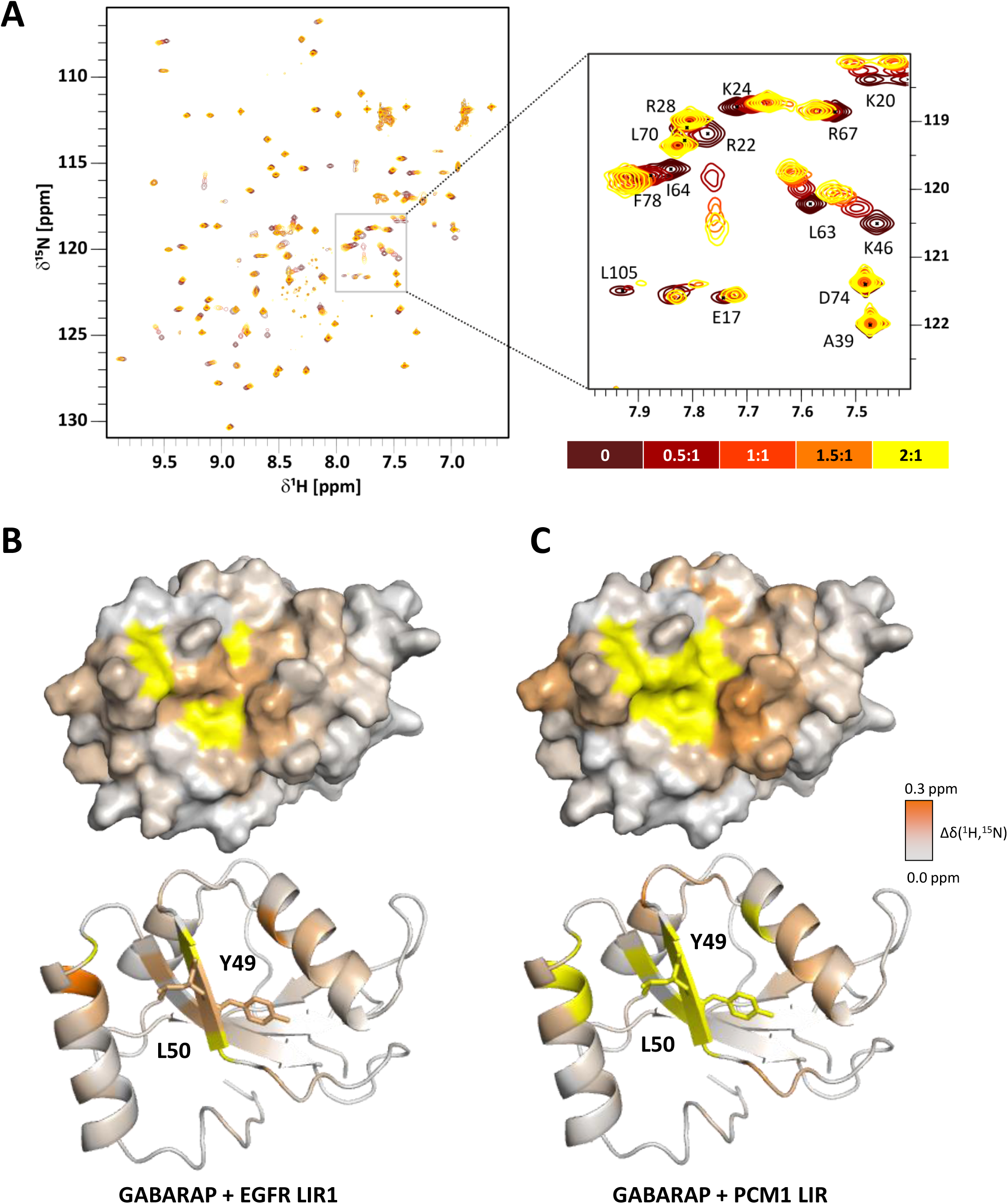
Interaction between GABARAP and LIR-peptides in solution. (**A**) 2D NMR spectrum (^1^H-^15^N-HSQC) of GABARAP incubated with increasing amounts of EGFR LIR1 and exemplary zoom-in of peaks representing affected residues. (**B, C**) Chemical shift perturbations during titration of ^15^N-GABARAP with LIR peptides from EGFR and PCM1 (2:1) mapped on the GABARAP structure (PDB ID: 1KOT) displayed as surface (top) and ribbon (bottom). Residues corresponding to disappearing HSQC peaks are presented in yellow.

### The interaction of EGFR LIR1 with GABARAP is independent of Y49A and L50A but is hindered by Pen8-*ortho*

To further verify the mode of binding, in particular the involvement of residues Y49 and L50 located in HP2 and HP1, respectively (see Fig. 4A), additional *in vitro* binding experiments were carried out. GABARAP_Y49A/L50A_ and GABARAPL1_Y49A/L50A_ were purified and subjected to BLI with immobilized PCM1 LIR and EGFR LIR1 respectively. In accordance with previous reports, PCM1 LIR binding was more than 20-fold reduced for GABARAP_Y49A/L50A_ (99.2 ± 6.7 µM) and GABARAPL1_Y49A/L50A_ (273.3 ± 21.8 µM) compared to the wildtype proteins (4.2 ± 0.6 and 4.9 ± 0.5 µM). In contrast, EGFR LIR1 binding was not influenced by the two mutations, neither for GABARAP/GABARAP_Y49A/L50A_ (52.0 ± 3.0 and 39.0 ± 2.6 µM), nor GABARAPL1/GABARAPL1_Y49A/L50A_ (29.7 ± 1.1 and 19.1 ± 1.2 µM, Fig. 4B). The ability of both GST-GABARAP and GST-GABARAP_Y49A/L50A_ to enrich endogenous EGFR from cell lysates, while within the same sample SQSTM1, described to depend on Y49 and L50 [49], showed a decrease with GST-GABARAP_Y49A/L50A_ (Fig. S6), notably supports the dispensability of these two GABARAP residues for EGFR (LIR1) binding. Finally, additional interaction studies were carried out using the stapled peptide Pen8-*ortho*, which has been described to bind to the GABARAP LDS with high affinity (K_D_: 14 nM) [29]. As expected, when preincubating GABARAP with a 1.5-fold molar excess of Pen8-*ortho,* 10-fold reduced binding to the immobilized EGFR LIR1 was observed (Fig. 4C). In conclusion, EGFR LIR1 appears to interact with GABARAP in a canonical, LDS-dependent manner, however, without relying on GABARAP residues Y49 and L50.

**Figure 4.**
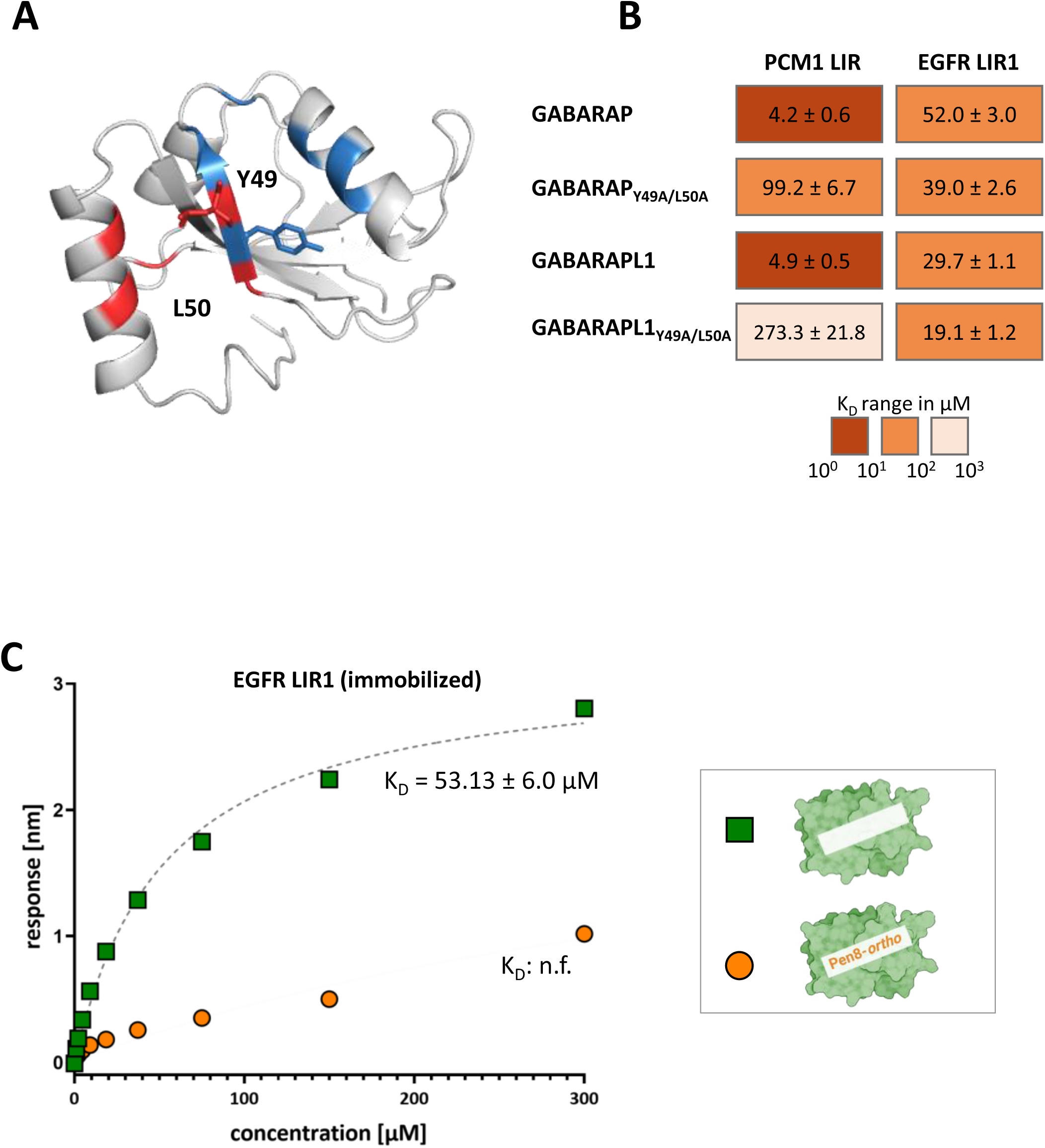
(**A**) Ribbon model of GABARAP with residues forming hydrophobic pockets HP1 and HP2 in red and blue, respectively. Residues Y49 and L50 are presented as sticks (**B**) K_D_ values [µM] of immobilized PCM1 and EGFR peptides with GABARAP/L1 and their Y49A/L50A variants as determined by biolayer interferometry. (**C**) Biolayer interferometry data for immobilized EGFR LIR1 peptide with either free GABARAP or GABARAP preincubated with the high-affinity binder Pen8-*ortho.* Effects of corresponding DMSO concentrations on binding were excluded (see Supplementary data for details). K_D_ values are shown with standard error calculated from non-linear regression. (n.f.: not fitable)

## Discussion

Receptor tyrosine kinase signalling, endocytic trafficking and autophagy are pathways important for cellular metabolism, which not only intersect at multiple stages, but also share several molecular components [50–53]. Following the observation of altered EGFR degradation and signaling, specifically in GABARAP single knockout cells [26], we sought to investigate a putative direct interaction between the hATG8 protein GABARAP and the EGFR. Interestingly, the core EGFR-LIR1 ‘FLPV’ possesses the same Θ_0_ and Γ_3_ residues as known hATG8 interactors preferentially binding the GABARAP subfamily, namely PCM1, ULK1, ULK2 and PIK3C3/VPS34 [20,21,54]. The X_1_ residue is occupied by a leucine, which shows van der Waals distance with hydrophobic amino acids of GABARAP in our X-ray structure. While being atypical for a GABARAP interaction motif (GIM), examples for non-selective hATG8 interactors with leucine in the X_1_ position, e.g. DVL2 have been described [21]. Notably, the X_2_ position, the least conserved residue of the consensus core LIR [55] and often not defining for GABARAP binding [56,57], is occupied by proline, a rather uncommon amino acid in this position. This P1088 does not appear to be involved in the interaction with GABARAP, possibly explaining the lower binding affinity of the EGFR LIR1 compared to related LIRs (e.g. in PCM1 and ULK1 [20]). Mutation studies have shown that proline residues in the X_2_ position are disruptive to the LIR-LDS binding [20,21], however in case of the PLEKHM1 LIR this effect was milder for GABARAP than LC3B [21]. Additionally, proline as X_2_ residues has been reported for functional LIRs of the valosin-containing protein (p97) as well as the pro-oxidant adaptor p66SHC [58,59]. Beyond the core, the EGFR LIR1 shares a proline as X_4_ residue (i.e. C-terminal to the core LIR) with ULK1, which has been suggested to inhibit LC3s binding [20], possibly explaining the lower EGFR LIR1 affinities for LC3A and LC3B. The N-terminal region preceding the core-LIR (X_-3_–X_-1_) has also frequently been reported to influence binding affinity and specificity, and acidic (D, E) as well as phorsphorylatable residues (S, T) appear to be common in these positions [55,60,61]. Indeed, N-terminal to the core LIR, the EGFR LIR1 has several acidic amino acids as well as serine and threonine, with X_-3_–X_-1_ being DDT, in fact, X_-3_ is located in proximity to H9 of GABARAP, possibly engaging in electrostatic interactions. Our investigation of the putative N-terminal phosphorylation sites, S1081 and T1085, revealed only slight increase in affinity. Interestingly, SCOC LIR phosphorylation did also only lead to slightly enhanced affinity for GABARAP [23], whereas in other cases, drastic effects of phosphorylation have been reported, e.g. ∼100-fold increase in affinity of LC3B to Nix upon phosphorylation of S34 and 35 in X_-1_ and X_-2_ [62].

EGFR internalization following EGF stimulation is regulated by parallel processes, namely clathrin-mediated endocytosis (CME), which itself can be categorized into AP2 dependent and independent CME, and non-clathrin endocytosis (NCE) which subsequently influence, degradation, recycling and signalling through multiple redundant and cooperative mechanisms [63–65]. Owing to the fact that neither these processes, nor the versatile roles of hATG8 proteins or possible compensatory effects resulting from GABARAP knockout are fully understood, pinpointing the molecular mechanism by which GABARAP influences the fate of the EGFR after EGF stimulation remains a challenge. While Dobner et al 2020 extensively discussed potential indirect mechanisms by which GABARAP could influence EGFR fate after stimulation [26], our data prompt us to also speculate on a direct interaction of the EGFR LIR1 with GABARAP. EGFR endocytosis and signaling is regulated by several binding sites within the CTD which are in proximity to the EGFR LIR1. This includes sites targeted by AP2 (positions 998–1001 (YLRA), 1034–1035 (LL) [65]), the palmitoyltransferase DHHC20 (cysteines at positions 1049, 1058, 1146 [66,67]) and the E3-ubiquitin ligase CBL (Y1069 [68,69]). Strikingly, the core LIR1 is located in the immediate vicinity of Y1092, a GRB2-binding site when phosphorylated [69,70]. Thus, LIR1 associated GABARAP could be suggested to hamper Y1092 phosphorylation itself and/or hamper GRB2 binding at pY1092, thus mitigating E3 ubiquitin-protein ligase CBL recruitment, EGFR ubiquitination and its lysosomal re-routing [69,70]. This provides a rationale for both the initially reduced pY1092 levels and the persistently higher levels of phosphorylated ERK observed in the presence of GABARAP compared to its absence [26]. Due to limited knowledge regarding the timeline of binding events following EGF-stimulation in presence and absence of GABARAP, obtaining solid experimental evidence remains challenging. Despite the scarcity of examples, LIR-LDS interactions between receptors and hATG8 have been proposed to be functionally relevant during diverse processes of receptor routing including their autophagic degradation (“signalophagy”) [71,72], anterograde trafficking [18] and secretion [17]. With our data, including the first X-ray structure of GABARAP with a surface receptor fragment, we propose the EGFR to be another of the yet few examples [73,74] of a receptor whose fate is influenced by an individual hATG8. Since receptor trafficking upon EGF treatment is altered in cells lacking GABARAP, direct hATG8-receptor interactions appear to not solely serve the autophagic degradation of RTKs in stressed cells, as shown for MET [71,72], but may alternatively play a general role during endosomal RTK transport in cells, possibly in a paralog-specific manner. Future studies designed to enrich our sparse understanding of RTK trafficking and associated LIR-dependencies will have to show whether this scenario proposed on the basis of *in vitro* binding data and the EGFR LIR1-GABARAP complex structure is indeed biologically relevant.

## Acknowledgement

This work was funded by the Deutsche Forschungsgemeinschaft (DFG, German Research Foundation)– Project-ID 267205415–SFB 1208 to D.W. Pen8-*ortho* was kindly supplied by Joshua Kritzer (Tufts University). We acknowledge the European Synchrotron Radiation Facility (ESRF) for provision of synchrotron radiation facilities and would like to thank the staff for assistance and support in using beamline ID30A-3.

## Author Contributions

Conceptualization: S.H., A.Ü.; investigation: A.Ü. (protein purification, BLI, X-ray, NMR; SI material), M.S. (NMR), O.H.W. (X-ray); formal analysis: A.Ü.; validation: S.H., A.Ü., O.H.W resources: D.W.; data curation: A.Ü., O.H.W.; visualization: A.Ü; writing & original draft preparation: A.Ü., S.H.; review and editing: all authors; supervision: S.H., O.H.W.; project administration: D.W.; funding acquisition: D.W.

## Abbreviations

AF2: AlphaFold 2
AP2: adaptor protein complex 2
BLI: biolayer interferometry
CASM: conjugation of ATG8 to single membranes
CBL: E3 ubiquitin-protein ligase CBL
CD36: platelet glycoprotein 4
CSP: chemical shift perturbation
CTD: carboxy-terminal domain
DHHC20: palmitoyltransferase ZDHHC20
DVL2: segment polarity protein dishevelled homolog
DVL-2 EGF: epidermal growth factor
EGFR: epidermal growth factor receptor
GABA_A_R: γ-aminobutyric acid type A receptor
GABARAP: γ-aminobutyric acid type A receptor-associated protein
GRB2: growth factor receptor-bound protein 2
GST: glutathione S-transferase
hATG8: human autophagy related protein 8
HSQC: heteronuclear single quantum coherence
LC3/MAP1LC3: microtubule-associated protein 1 light chain 3
LDELS: LC3-dependent extracellular vesicle loading and secretion
LDS: LIR docking site
LIR: LC3-interacting region
MAPK/ERK: mitogen-activated protein kinase/extracellular-signal regulated kinases
PCM1: pericentriolar material 1 protein
p66SHC: p66 isoform of Src homology 2 domain-containing transforming protein 1 (SHC1)
PIK3C3/VPS34: phosphatidylinositol 3-kinase catalytic subunit type 3/
SCOC: short coiled-coil protein
SQSTM1: sequestosome-1
TLR4: toll-like receptor 4
TREM2: triggering receptor expressed on myeloid cells 2
ULK1: unc-51-like kinase 1

