## Supporting Information for "EGFR meets hATG8s – Biophysical and structural insights supporting a unique role of GABARAP during receptor trafficking"

### SI Figures

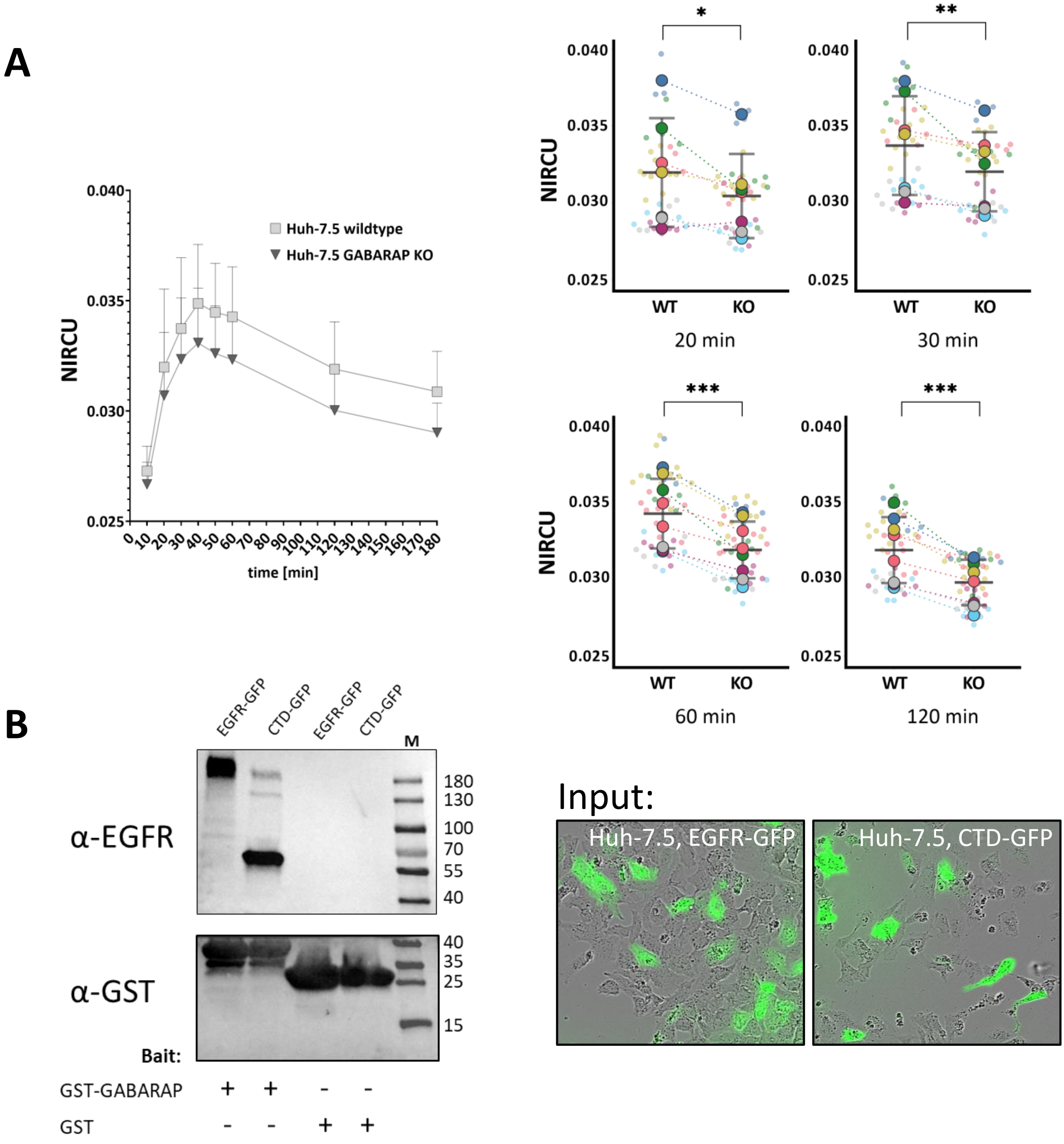

**Figure S1.** (**A**) EGF-647 levels in Huh-7.5 wildtype (WT) and GABARAP KO (KO) cells over time measured in near infrared calibrated units (NIRCU) **B**) Huh-7.5 cells were transfected with plasmids encoding EGFR-GFP and CTD-GFP respectively. Lysates were used for affinity enrichment with GST-GABARAP. GST only was used as control. The respective western blot was stained with antibodies against EGFR and GST.

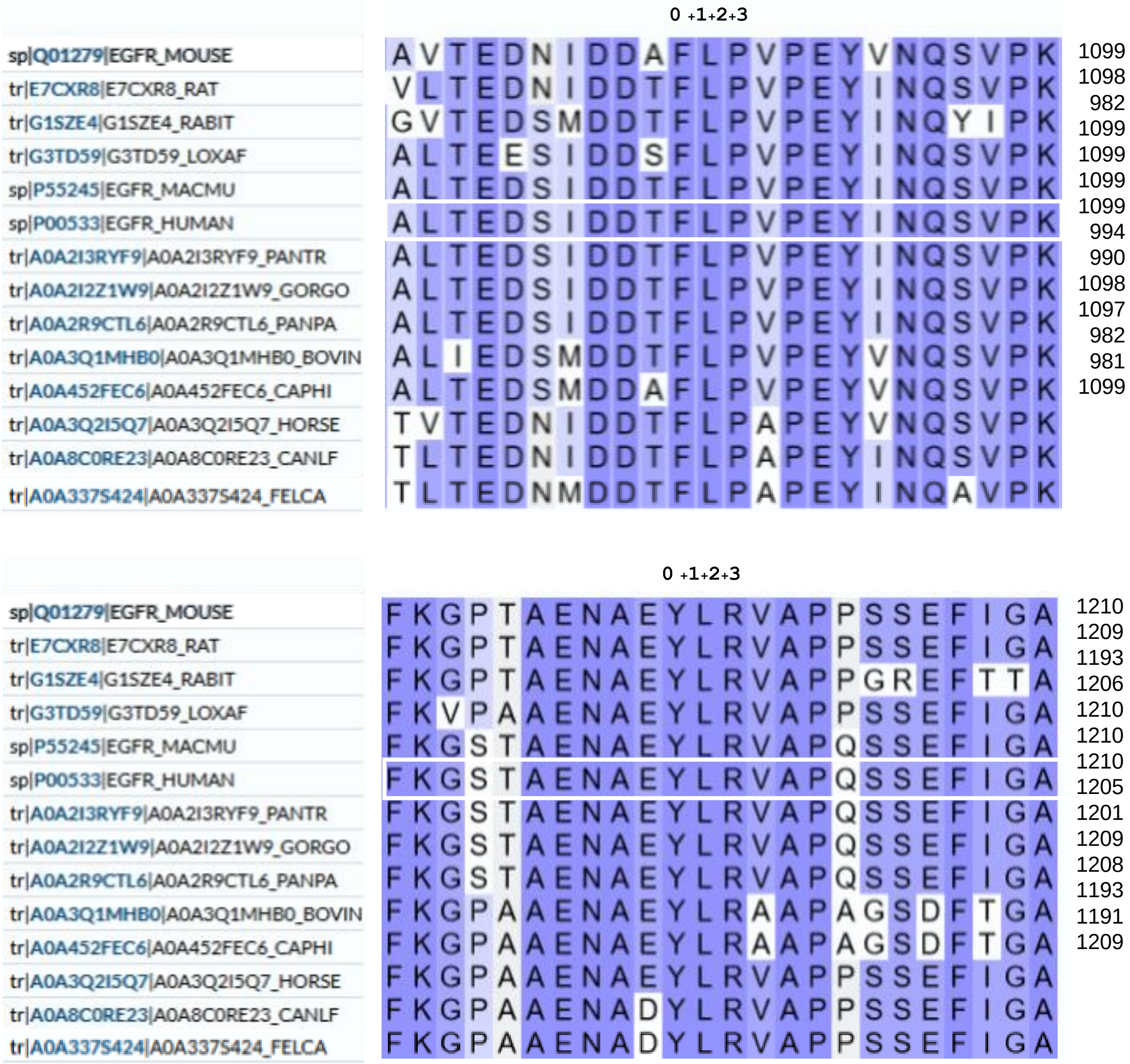

**Figure S2.** Conservation of LIR1 and LIR2 surrounding regions within mammals. The multiple sequence alignment for the selected EGFR protein sequences (full-length) was obtained using the Align function of the UniProt Consortium [4] according to described protocols [5]. The human EGFR sequence is highlighted by a white box. Only the two relevant sequence segments (top: LIR1, bottom: LIR2) are shown with the corresponding LIR-core positions numbered from 0 to +3. Interestingly, if divergent, position +3 in both core LIRs appears to be preferentially occupied by alanine.

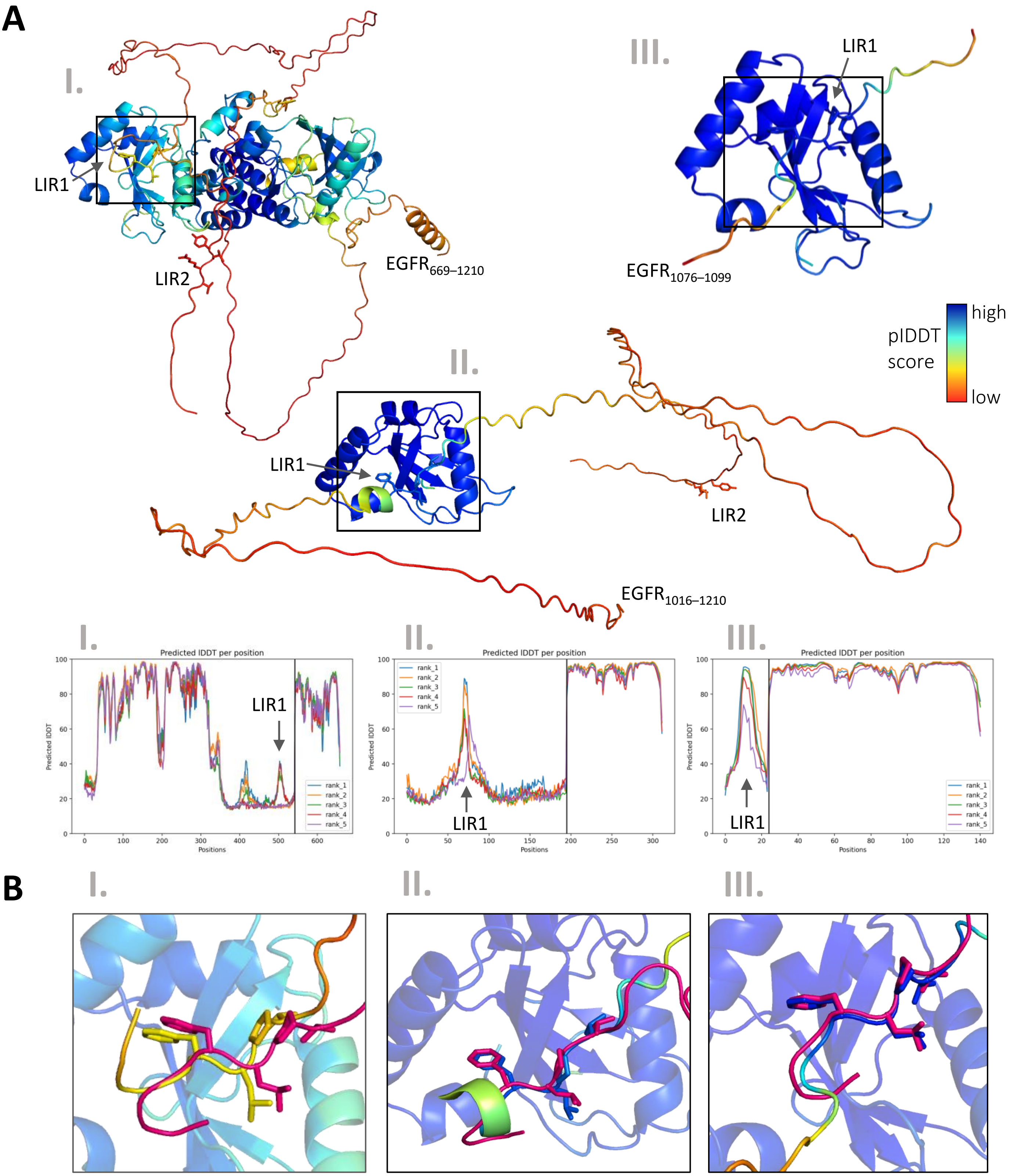

**Figure S3.** Prediction of GABARAP binding site selection within the EGFR cytoplasmic region (**A**). AF2-multimer prediction of complex structures of GABARAP and the EGFR cytoplasmic domain (I.) or fragments thereof (II. and III.). Models (colored according their respective lDDT scores) along with their predicted lDDT plots are shown. As both LIR1 and LIR2 are located within intrinsically disordered protein regions of EGFR, an AF2-related bias (lower selection probability for LIRs in more structured regions [6]) for either of them is not expected. PTMs are neglected as they cannot be processed by AF2. Although a canonical LIR2-GABARAP LDS interaction was predicted for the short EGFR fragment (1190–1210) as input, LIR2 was not selected with any of the larger EGFR CTD fragments tested, their LIR1-mutated (FLPV to ALPA) versions or when offering two GABARAP molecules as input (not shown). (**B**) Superposition of the X-ray structure (8S1M) with AF2 rank 1 models of GABARAP in complex with LIR1 in the context of the entire cytoplasmic domain (I.), the EGFR fragment 1016–1210 (II.) and the 24mer peptide (III.), used for BLI. Only respective core LIR residues are depicted with those from 8S1M colored in pink. IDDT: Local Distance Difference Test

| | GABARAP | GABARAPL1 | GABARAPL2 | LC3A | LC3B |
|---|---|---|---|---|---|
| pY1092 EGFR LIR1 | 46.23 ± 4.45 | 19.5 ± 0.6 | 98.0 ± 10.8 | 135.7 ± 6.9 | 337.9 ± 20.9 |
| pY1197 EGFR LIR2 | 302.31 ± 26.61 | 30.9 ± 1.4 | 199.2 ± 15.8 | 229.7 ± 12.3 | 511.2 ± 39.0 |

$K_D$ range in µM

$10^0$ $10^1$ $10^2$ >300

**Figure S4.** $K_D$ values [µM] with standard error calculated from non-linear regression of pY1092 EGFR-LIR1 and pY1197 EGFR-LIR2 (phosphorylated core LIR tyrosine) and human ATG8 paralogs.

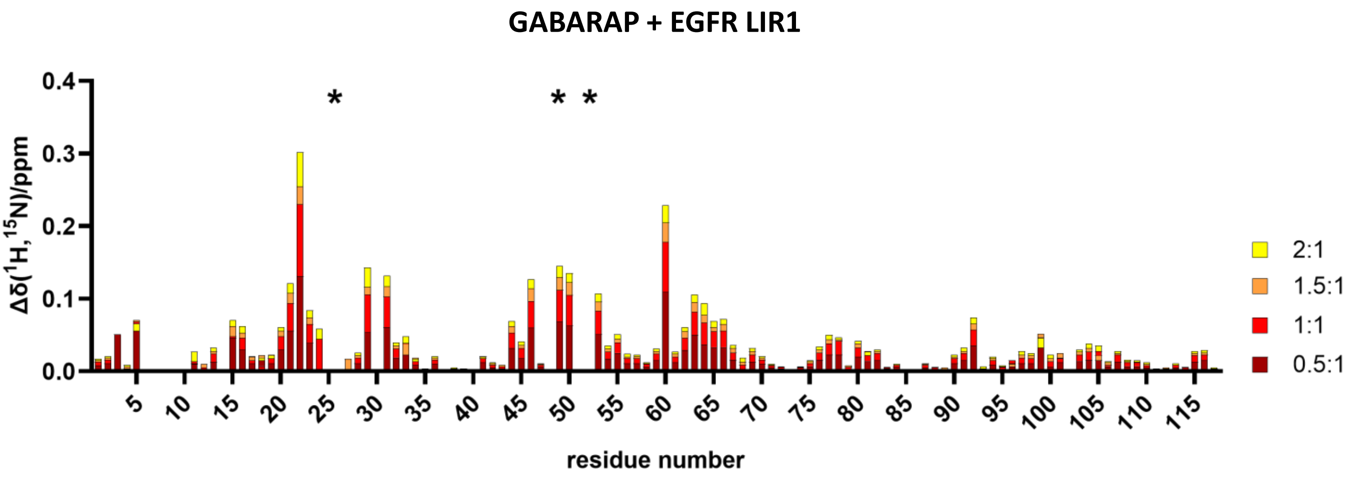

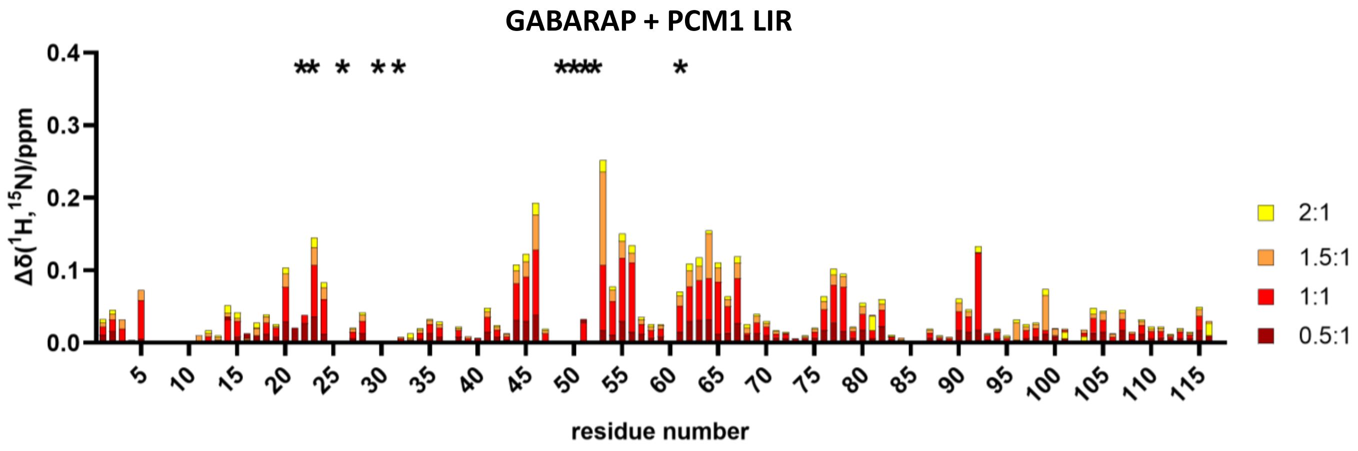

**Figure S5.** CSP for backbone amide resonances of GABARAP incubated with the indicated stoichiometric ratios of EGFR LIR1 (top) and PCM1 LIR (bottom) peptides. Asterisks represent residues affected by strong line broadening beyond detection - Y25, K48 and V51 in case of EGFR LIR1 and I21, R22, Y24, K48, Y49, L50, V51, F60 in case of PCM1 LIR.

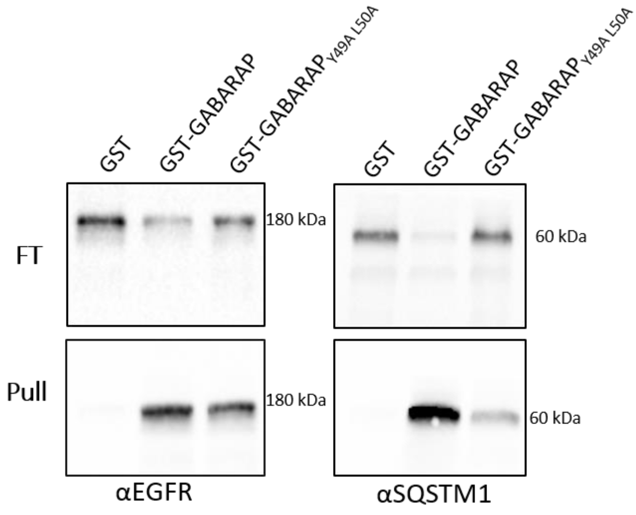

**Figure S6.** Affinity enrichment of endogenous EGFR (left) and SQSTM1 (right) from Huh-7.5 cells with GST-GABARAP and GST-GABARAPY$_{49A/L50A}$ affinity purified from bacterial lysate.

### SI Tables

**Supplementary Table 1: Peptides used for BLI affinity measurements** (phosphorylation is indicated by p).

| Peptide | Sequence |
|---|---|
| EGFR LIR1 | Biotin-Ahx-ALTEDSIDDTFLPVPEYINQSVPK-$NH_2$ |
| (pS)EGFR LIR1 | Biotin-Ahx-ALTED(pS)IDDTFLPVPEYINQSVPK-$NH_2$ |
| (pT)EGFR LIR1 | Biotin-Ahx-ALTEDSIDD(pT)FLPVPEYINQSVPK-$NH_2$ |
| (pSpT)EGFR LIR1 | Biotin-Ahx-ALTED(pS)IDD(pT)FLPVPEYINQSVPK-$NH_2$ |
| (pY)EGFR LIR1 | Biotin-Ahx-ALTEDSIDDTFLPVPE(pY)INQSVPK-$NH_2$ |
| EGFR LIR2 | Biotin-Ahx-STAENAEYLRVAPQSSEFIGA-$NH_2$ |
| (pY)EGFR LIR2 | Biotin-Ahx-STAENAEYLRVAPQSSEFIGA-$NH_2$ |
| PCM1 LIR (1954–1968) | Biotin-Ahx-SQKSDEEDFVKVEDLPLKLTI-$NH_2$ |

Ahx: Aminohexanoic acid linker

**Supplementary Table 2: RNA expression profile of hATG8.** Data are extracted from the Cell Lines section of the Human Protein Atlas (https://www.proteinatlas.org/humanproteome/cell+line; v23).
nTPM: normalized transcripts per million.

| hATG8 | nTPM | |
|---|---|---|
| | HEK293 | Huh7 |
| GABARAP | 445.0 | 496.4 |
| GABARAPL1 | 28.0 | 43.3 |
| GABARAPL2 | 195.8 | 112.1 |
| LC3A | 1.9 | 13.4 |
| LC3B | 76.0 | 79.2 |
| LC3B2 | 1.8 | 2.1 |
| LC3C | 1.0 | 0 |

**Supplementary Table 3: Data collection and refinement statistics.**

| | EGFR LIR1-GABARAP |
|---|---|
| **Data collection** | |
| Space group | I23 |
| Cell dimensions | |
| a, b, c (Å) | 101.28, 101.28, 101.28 |
| α, β, γ (°) | 90, 90, 90 |
| Resolution (Å) | 41.35–2.05 (2.10–2.05) * |
| $R_{measure}$ | 0.076 (3.308) |
| CC1/2 | 1 (0.365) |
| Mean I/σ(I) | 21.91 (1.08) |
| Completeness (%) | 99.95 (100.00) |
| Multiplicity | 20.09 (20.10) |
| **Refinement** | |
| Resolution (Å) | 41.35–2.05 |
| No. reflections | 11007 (1081) |
| $R_{work}$ / $R_{free}$ | 0.2019/ 0.2286 |
| No. atoms | 1134 |
| Protein | 1101 |
| Ligand/ion | 7 |
| Water | 26 |
| Mean B-factors ($Å^2$) | 72.05 |
| Protein | 72.13 |
| Ligand/ion | 84.31 |
| Water | 65.20 |
| R.m.s. deviations | |
| Bond lengths (Å) | 0.005 |
| Bond angles (°) | 0.67 |

*Values in parentheses are for the highest-resolution shell.

### SI Methods

#### Cell culture

Human hepatoma Huh-7.5 wildtype and GABARAP KO cells [1] were grown in Dulbecco's Modified Eagle Medium (DMEM) high glucose (Sigma Aldrich, D5796), supplemented with 10% heat inactivated fetal bovine serum (Sigma Aldrich, F9665) and 1% penicillin/streptomycin (Sigma Aldrich, P4333) at 37°C and 5% $CO_2$. Cells were passaged at approximately 80% confluency. For transfection, cells were grown in 6-well plates to at least 70% confluency and transfected with 3 µg plasmid DNA per well, using 9 µl Lipofectamine 2000 (Thermo Fisher Scientific, 11668019).

#### Continuous live-cell microscopy

For continuously monitoring the levels of EGF-Alexa647 in Huh-7.5 wildtype and GABARAP KO cells, real-time live cell microscopy (Incucyte SX5, Sartorius) was applied. The day prior to the experiment, 10,000 cells per well were seeded into 96 well plates. The following day, cells were cooled on ice for 10 min and 40 ng/ml EGF Alexa Fluor 647 (EGF-647; Thermo Fisher Scientific, E35351) in cold medium was added to the cells, which were incubated for one hour at 4°C on ice protected from light to allow ligand binding to the receptor. Afterwards, the medium was exchanged to remove free EGF-647 and cells were transferred into the Incucyte SX5, where images were recorded every 10 min for one hour and subsequently every hour using the 'adherent cell-by-cell' mode and the Phase and near infrared (NIR) channels. For analysis, the Incucyte cell-by-cell software (version 2021C; Sartorius) was applied to analyze the fluorescence intensity per cell over time. Cells with exceptionally high fluorescence intensity (above 145 NIR calibrated units (NIRCU), less than 2% of all cells) were excluded.

#### Affinity enrichment and immunoblotting

Huh-7.5 (either untransfected or transfected with pEGFR-GFP/pCTD-GFP, coding for full-length EGFR or for residues 971 to 1210 comprising the regulatory C-terminal domain of the receptor) cells were harvested by trypsination, washed once in PBS and lysed with NP40 buffer (20 mM Tris HCl, 200 mM NaCl, 1 mM EDTA, 0.5% NP40, 1 mM phenylmethylsulfonylfluoride (PMSF) and 1x Halt protease and phosphatase inhibitor [Thermo Fisher Scientific, 78442]) by incubation on ice for 30 min and vigorous pipetting every 10 min. Insoluble cell parts were sedimented by centrifugation for 10 min at 17,000 x g. GSH Sepharose beads (GE Healthcare, 17075605) were loaded with respective proteins by incubation with bacterial lysate containing GST-GABARAP, GST-GABARAP$_{Y49A/L50A}$ (refer to main text for details on expression) or purified GST, respectively and thoroughly washed. The cell lysates were precleared (GST loaded beads for 2 hours at 4°C) and subsequently incubated overnight at 4°C with GST-GABARAP, GST-GABARAP$_{Y49A/L50A}$ or GST loaded GSH beads, respectively. Beads were washed four times with cell lysis buffer, combined with Lämmli buffer and heated to 95°C for 10 minutes. Subsequently, samples were separated by SDS-PAGE using precast stain-free gels (Bio-Rad Laboratories, 4568124), transferred to 0.2 µM PVDF membranes (Bio-Rad

Laboratories, 1704156) and blocked in 5% BSA (AppliChem, A1391) in TBS-T (136 mM NaCl, 2.7 mM KCl, 24.7 mM Tris-HCl, pH 7.4, 0.05% Tween-20 [Applichem, A4974]) for 1 h. Membrane was incubated with primary antibody against EGFR (Cell Signaling Technology, 4267, rabbit, 1:1000), SQSTM1 (BD Bioscience, 610832, mouse, 1:1000) and GST (Novus biologicals NB600-388A, 1:1000, HRP conjugated) overnight at 4°C, washed 3 times with TBS-T, incubated with secondary antibodies (anti-rabbit IgG/HRP [Dako, P0448, 1:5000] & anti-mouse IgG/HRP [Dako, P0260, 1:1000] for 1 h at room temperature and again washed thrice with TBS-T. GST-related signals were visualized with ECL Select Western Blotting Detection Reagent (cytiva, RPN2235) or Clarity Western Substrate (Biorad, #1705061).

##### Alphafold predictions

Complex structures of GABARAP and the EGFR cytoplasmic domain (or fragments thereof) were investigated *in silico* with AlphaFold2 [2], using the ColabFold implementation [3-6]. For each subject, five models were generated without inclusion of template information, using multiple sequence alignment mode mmseqs2_uniref_env, model type alphafold2_multimer_v3, a maximum of 20 recycle steps, and recycling controlled by recycle_early_stop_tolerance: auto. Models for GABARAP complexed with EGFR 1016–1210 and EGFR 1076–1099 were additionally subjected to relaxation using Amber as implemented in the ColabFold pipeline. The highest-ranked model (according to predicted lDDT statistics) without steric clashes was used for further evaluation. Figures were created using the PyMOL Molecular Graphics System, Version 2.7 Schrödinger, LLC.

#### Fig. 1B Bio-Layer Interferometry (BLI) affinity measurements of EGFR derived peptides with hATG8 paralogs

| | EGFR LIR1 |
|---|---|
| GABARAP [μM] | Response |
| 0 | -0.0081 |
| 1.125 | 0.1081 |
| 2.25 | 0.1894 |
| 4.55 | 0.3376 |
| 9.123 | 0.5630 |
| 18.62 | 0.8780 |
| 37.25 | 1.2865 |
| 75 | 1.74951 |
| 150 | 2.24111 |
| 300 | 2.80478 |

| **Best-fit values** | |
|---|---|
| $B_{max}$ | 3.156 |
| $K_D$ | 53.13 |
| **Standard Error** | |
| $B_{max}$ | 0.123 |
| $K_D$ | 6.011 |
| **95% Confidence Intervals** | |
| $B_{max}$ | 2.881 to 3.498 |
| $K_D$ | 40.21 to 70.49 |
| **Goodness of Fit** | |
| R square | 0.993 |

| | EGFR LIR2 |
|---|---|
| GABARAP [μM] | Response |
| 0 | -0.0099 |
| 1.125 | 0.0046 |
| 2.25 | 0.0128 |
| 4.55 | 0.0390 |
| 9.123 | 0.0682 |
| 18.62 | 0.1287 |
| 37.25 | 0.2299 |
| 75 | 0.38441 |
| 150 | 0.62864 |
| 300 | 0.98236 |

| **Best-fit values** | |
|---|---|
| $B_{max}$ | 1.963 |
| $K_D$ | 304.3 |
| **Standard Error** | |
| $B_{max}$ | 0.099 |
| $K_D$ | 25.386 |
| $B_{max}$ | 1.755 to 2.231 |
| $K_D$ | 251.8 to 374.3 |
| **Goodness of Fit** | |
| R square | 0.9987 |

| | PCM1 LIR |
|---|---|
| GABARAP [μM] | Response |
| 0 | -0.0090 |
| 0.19 | 0.2321 |
| 0.39 | 0.4139 |
| 0.78 | 0.6859 |
| 1.56 | 1.0623 |
| 3.125 | 1.5184 |
| 6.25 | 2.0116 |
| 12.5 | 2.45342 |
| 25 | 2.86824 |
| 50 | 3.28205 |

| **Best-fit values** | |
|---|---|
| $B_{max}$ | 3.352 |
| $K_D$ | 3.722 |
| **Standard Error** | |
| $B_{max}$ | 0.095 |
| $K_D$ | 0.317 |
| **95% Confidence Intervals** | |
| $B_{max}$ | 3.133 to 3.590 |
| $K_D$ | 2.909 to 4.755 |
| **Goodness of Fit** | |
| R square | 0.9931 |

| | EGFR LIR1 |
|---|---|
| GABARAPL1 [μM] | Response |
| 0 | -0.0003 |
| 1.125 | 0.1310 |
| 2.25 | 0.2229 |
| 4.55 | 0.4023 |
| 9.123 | 0.6381 |
| 18.62 | 0.9541 |
| 37.25 | 1.3168 |
| 75 | 1.64362 |
| 150 | 1.88425 |
| 300 | 1.93841 |

| **Best-fit values** | |
|---|---|
| $B_{max}$ | 21.63 |
| $K_D$ | 2.108 |
| **Standard Error** | |
| $B_{max}$ | 0.023 |
| $K_D$ | 0.84 |
| **95% Confidence Intervals** | |
| $B_{max}$ | 2.055 to 2.162 |
| $K_D$ | 19.76 to 23.67 |
| **Goodness of Fit** | |
| R square | 0.999 |

| | EGFR LIR2 |
|---|---|
| GABARAPL1 [μM] | Response |
| 0 | -0.0043 |
| 1.125 | 0.0160 |
| 2.25 | 0.0236 |
| 4.55 | 0.0645 |
| 9.123 | 0.1124 |
| 18.62 | 0.1888 |
| 37.25 | 0.2955 |
| 75 | 0.41547 |
| 150 | 0.52585 |
| 300 | 0.57995 |

| **Best-fit values** | |
|---|---|
| $B_{max}$ | 0.6809 |
| $K_D$ | 48.04 |
| **Standard Error** | |
| $B_{max}$ | 0.008 |
| $K_D$ | 1.619 |
| **95% Confidence Intervals** | |
| $B_{max}$ | 0.6633 to 0.6993 |
| $K_D$ | 44.48 to 51.90 |
| **Goodness of Fit** | |
| R square | 0.9994 |

| | PCM1 LIR |
|---|---|
| GABARAPL1 [μM] | Response |
| 0 | 0.0135 |
| 0.147 | 0.2371 |
| 0.293 | 0.3916 |
| 0.586 | 0.6283 |
| 1.172 | 0.9687 |
| 2.344 | 1.4215 |
| 4.688 | 1.8478 |
| 9.376 | 2.22826 |
| 18.75 | 2.6018 |

| **Best-fit values** | |
|---|---|
| $B_{max}$ | 2.828 |
| $K_D$ | 2.258 |
| **Standard Error** | |
| $B_{max}$ | 0.067 |
| $K_D$ | 0.17 |
| **95% Confidence Intervals** | |
| $B_{max}$ | 2.670 to 2.999 |
| $K_D$ | 1.876 to 2.718 |
| **Goodness of Fit** | |
| R square | 0.9966 |

| | EGFR LIR1 |
|---|---|
| GABARAPL2 [μM] | Response |
| 0 | 0.0021 |
| 1.777 | 0.0336 |
| 3.554 | 0.0493 |
| 7.109 | 0.0794 |
| 14.21 | 0.1303 |
| 28.44 | 0.2150 |
| 56.88 | 0.3752 |
| 113.8 | 0.57488 |
| 227.5 | 0.78398 |
| 455 | 0.87138 |

| **Best-fit values** | |
|---|---|
| $B_{max}$ | 1.098 |
| $K_D$ | 104.8 |
| **Standard Error** | |
| $B_{max}$ | 0.031 |
| $K_D$ | 7.712 |
| **95% Confidence Intervals** | |
| $B_{max}$ | 1.032 to 1.171 |
| $K_D$ | 88.80 to 123.9 |
| **Goodness of Fit** | |
| R square | 0.9974 |

| | EGFR LIR2 |
|---|---|
| GABARAPL2 [μM] | Response |
| 0 | -0.0046 |
| 1.777 | 0.0101 |
| 3.554 | 0.0114 |
| 7.109 | 0.0196 |
| 14.21 | 0.0268 |
| 28.44 | 0.0428 |
| 56.88 | 0.0785 |
| 113.8 | 0.12363 |
| 227.5 | 0.18022 |
| 455 | 0.21482 |

| **Best-fit values** | |
|---|---|
| $B_{max}$ | 0.2871 |
| $K_D$ | 146.2 |
| **Standard Error** | |
| $B_{max}$ | 0.011 |
| $K_D$ | 13.158 |
| **95% Confidence Intervals** | |
| $B_{max}$ | 0.2640 to 0.3141 |
| $K_D$ | 119.2 to 180.3 |
| **Goodness of Fit** | |
| R square | 0.9965 |

| | PCM1 LIR |
|---|---|
| GABARAPL2 [μM] | Response |
| 0 | -0.0113 |
| 0.888671875 | 0.2187 |
| 1.77734375 | 0.3374 |
| 3.5546875 | 0.5476 |
| 7.109375 | 0.8827 |
| 14.21875 | 1.2250 |
| 28.4375 | 1.6078 |
| 56.875 | 2.00653 |
| 113.75 | 2.38051 |

| **Best-fit values** | |
|---|---|
| $B_{max}$ | 2.595 |
| $K_D$ | 15.01 |
| **Standard Error** | |
| $B_{max}$ | 0.085 |
| $K_D$ | 1.514 |
| **95% Confidence Intervals** | |
| $B_{max}$ | 2.397 to 2.818 |
| $K_D$ | 11.66 to 19.36 |
| **Goodness of Fit** | |
| R square | 0.994 |

| | EGFR LIR1 |
|---|---|
| LC3A [μM] | Response |
| 0 | -0.0505 |
| 2.929 | -0.0093 |
| 5.859 | 0.0309 |
| 11.72 | 0.1450 |
| 23.44 | 0.2486 |
| 46.88 | 0.4744 |
| 93.75 | 0.8254 |
| 187.5 | 1.22173 |
| 375 | 1.67254 |
| 750 | 2.12348 |

| **Best-fit values** | |
|---|---|
| $B_{max}$ | 2.753 |
| $K_D$ | 231.2 |
| **Standard Error** | |
| $B_{max}$ | 0.071 |
| $K_D$ | 14.341 |
| **95% Confidence Intervals** | |
| $B_{max}$ | 2.597 to 2.927 |
| $K_D$ | 200.4 to 267.6 |
| **Goodness of Fit** | |
| R square | 0.9984 |

| | EGFR LIR2 |
|---|---|
| LC3A [μM] | Response |
| 0 | -0.0151 |
| 2.929 | -0.0086 |
| 5.859 | 0.0079 |
| 11.72 | 0.0343 |
| 23.44 | 0.0442 |
| 46.88 | 0.0917 |
| 93.75 | 0.1799 |
| 187.5 | 0.29706 |
| 375 | 0.48793 |
| 750 | 0.72264 |

| **Best-fit values** | |
|---|---|
| $B_{max}$ | 1.344 |
| $K_D$ | 648.6 |
| **Standard Error** | |
| $B_{max}$ | 0.06 |
| $K_D$ | 50.695 |
| **95% Confidence Intervals** | |
| $B_{max}$ | 1.217 to 1.502 |
| $K_D$ | 544.0 to 783.5 |
| **Goodness of Fit** | |
| R square | 0.9987 |

| | PCM1 LIR |
|---|---|
| LC3A [μM] | Response |
| 0 | -0.0373 |
| 2.929 | 0.0923 |
| 5.859 | 0.2182 |
| 11.72 | 0.4316 |
| 23.44 | 0.7120 |
| 46.88 | 1.1687 |
| 93.75 | 1.7259 |
| 187.5 | 2.32909 |
| 375 | 2.92617 |
| 750 | 3.49212 |

| **Best-fit values** | |
|---|---|
| $B_{max}$ | 3.936 |
| $K_D$ | 118 |
| **Standard Error** | |
| $B_{max}$ | 0.089 |
| $K_D$ | 7.839 |
| **95% Confidence Intervals** | |
| $B_{max}$ | 3.733 to 4.156 |
| $K_D$ | 100.7 to 138.5 |
| **Goodness of Fit** | |
| R square | 0.9975 |

| | EGFR LIR1 |
|---|---|
| LC3B [μM] | Response |
| 0 | -0.0039 |
| 3.906 | 0.0017 |
| 7.812 | 0.0307 |
| 15.63 | 0.0824 |
| 31.25 | 0.1375 |
| 62.5 | 0.2392 |
| 125 | 0.4130 |
| 250 | 0.64423 |
| 500 | 0.96087 |
| 1000 | 1.27008 |

| **Best-fit values** | |
|---|---|
| $B_{max}$ | 1.811 |
| $K_D$ | 434.6 |
| **Standard Error** | |
| $B_{max}$ | 0.04 |
| $K_D$ | 20.851 |
| **95% Confidence Intervals** | |
| $B_{max}$ | 1.721 to 1.910 |
| $K_D$ | 388.8 to 487.1 |
| **Goodness of Fit** | |
| R square | 0.9992 |

| | EGFR LIR2 |
|---|---|
| LC3B [μM] | Response |
| 0 | -0.0108 |
| 3.906 | -0.0245 |
| 7.812 | -0.0119 |
| 15.63 | 0.0104 |
| 31.25 | 0.0113 |
| 62.5 | 0.0228 |
| 125 | 0.0556 |
| 250 | 0.09588 |
| 500 | 0.18218 |
| 1000 | 0.27646 |

| **Best-fit values** | |
|---|---|
| $B_{max}$ | 0.6859 |
| $K_D$ | 1461 |
| **Standard Error** | |
| $B_{max}$ | 0.159 |
| $K_D$ | 504.122 |
| **95% Confidence Intervals** | |
| $B_{max}$ | 0.4558 to 1.550 |
| $K_D$ | 755.3 to 4285 |
| **Goodness of Fit** | |
| R square | 0.9864 |

| | PCM1 LIR |
|---|---|
| LC3B [μM] | Response |
| 0 | -0.00927 |
| 3.906 | -0.00792 |
| 7.812 | 0.03144 |
| 15.63 | 0.10506 |
| 31.25 | 0.1975 |
| 62.5 | 0.36936 |
| 125 | 0.62677 |
| 250 | 0.97778 |
| 500 | 1.39609 |
| 1000 | 1.7934 |

| **Best-fit values** | |
|---|---|
| $B_{max}$ | 2.444 |
| $K_D$ | 368.7 |
| **Standard Error** | |
| $B_{max}$ | 0.042 |
| $K_D$ | 14.514 |
| **95% Confidence Intervals** | |
| $B_{max}$ | 2.350 to 2.546 |
| $K_D$ | 336.7 to 404.4 |
| **Goodness of Fit** | |
| R square | 0.9994 |

#### Fig. 1C Bio-Layer Interferometry (BLI) affinity measurements of EGFR derived phosphopeptides with GABARAP

| | EGFR LIR1 |
|---|---|
| GABARAP [μM] | Response |
| 0 | 0.0012 |
| 1.125 | 0.0874 |
| 2.25 | 0.1527 |
| 4.55 | 0.2731 |
| 9.123 | 0.4764 |
| 18.62 | 0.7470 |
| 37.25 | 1.1314 |
| 75 | 1.59117 |
| 150 | 2.07138 |
| 300 | 2.52601 |

| **Best-fit values** | |
|---|---|
| $B_{max}$ | 2.948 |
| $K_D$ | 58.76 |
| **Standard Error** | |
| $B_{max}$ | 0.081 |
| $K_D$ | 4.437 |
| **95% Confidence Intervals** | |
| $B_{max}$ | 2.765 to 3.152 |
| $K_D$ | 49.02 to 70.59 |
| **Goodness of Fit** | |
| R square | 0.997 |

| | (pY)EGFR LIR1 |
|---|---|
| GABARAP [μM] | Response |
| 0 | -0.0003 |
| 1.125 | 0.0862 |
| 2.25 | 0.1538 |
| 4.55 | 0.2801 |
| 9.123 | 0.4887 |
| 18.62 | 0.7849 |
| 37.25 | 1.1709 |
| 75 | 1.63648 |
| 150 | 2.10076 |
| 300 | 2.51907 |

| **Best-fit values** | |
|---|---|
| $B_{max}$ | 2.907 |
| $K_D$ | 53.88 |
| **Standard Error** | |
| $B_{max}$ | 0.066 |
| $K_D$ | 3.457 |
| **95% Confidence Intervals** | |
| $B_{max}$ | 2.756 to 3.072 |
| $K_D$ | 46.18 to 62.94 |
| **Goodness of Fit** | |
| R square | 0.9977 |

| | (pS)EGFR LIR1 |
|---|---|
| GABARAP [μM] | Response |
| 0 | -0.0054 |
| 1.125 | 0.1869 |
| 2.25 | 0.3314 |
| 4.55 | 0.5827 |
| 9.123 | 0.9572 |
| 18.62 | 1.4027 |
| 37.25 | 1.9270 |
| 75 | 2.4847 |
| 150 | 3.02065 |
| 300 | 3.49246 |

| **Best-fit values** | |
|---|---|
| $B_{max}$ | 3.715 |
| $K_D$ | 31.72 |
| **Standard Error** | |
| $B_{max}$ | 0.105 |
| $K_D$ | 2.918 |
| **95% Confidence Intervals** | |
| $B_{max}$ | 3.474 to 3.981 |
| $K_D$ | 25.31 to 39.75 |
| **Goodness of Fit** | |
| R square | 0.9946 |

| | (pT)EGFR LIR1 |
|---|---|
| GABARAP [μM] | Response |
| 0 | -0.0060 |
| 1.125 | 0.1795 |
| 2.25 | 0.3237 |
| 4.55 | 0.5647 |
| 9.123 | 0.9314 |
| 18.62 | 1.3561 |
| 37.25 | 1.8759 |
| 75 | 2.43907 |
| 150 | 2.95936 |
| 300 | 3.42416 |

| **Best-fit values** | |
|---|---|
| $B_{max}$ | 3.655 |
| $K_D$ | 32.4 |
| **Standard Error** | |
| $B_{max}$ | 0.102 |
| $K_D$ | 2.935 |
| **95% Confidence Intervals** | |
| $B_{max}$ | 3.420 to 3.914 |
| $K_D$ | 25.95 to 40.45 |
| **Goodness of Fit** | |
| R square | 0.9948 |

| | (pSpT)EGFR LIR1 |
|---|---|
| GABARAP [μM] | Response |
| 0 | -0.0085 |
| 1.125 | 0.2720 |
| 2.25 | 0.4753 |
| 4.55 | 0.7964 |
| 9.123 | 1.2234 |
| 18.62 | 1.6804 |
| 37.25 | 2.1954 |
| 75 | 2.72107 |
| 150 | 3.21036 |
| 300 | 3.61812 |

| **Best-fit values** | |
|---|---|
| $B_{max}$ | 3.696 |
| $K_D$ | 21.73 |
| **Standard Error** | |
| $B_{max}$ | 0.114 |
| $K_D$ | 2.38 |
| **95% Confidence Intervals** | |
| $B_{max}$ | 3.431 to 3.986 |
| $K_D$ | 16.53 to 28.51 |
| **Goodness of Fit** | |
| R square | 0.9915 |

#### Fig. 4B Bio-Layer Interferometry (BLI) affinity measurements of EGFR LIR1 and PCM1 LIR GABARAP/L1 and Y49A L50A

| | EGFR LIR1 |
|---|---|
| GABARAP [μM] | Response |
| 0 | -0.0015 |
| 0.781 | 0.0462 |
| 1.562 | 0.0724 |
| 3.125 | 0.1310 |
| 6.25 | 0.2293 |
| 12.5 | 0.3748 |
| 25 | 0.6059 |
| 50 | 0.88171 |
| 100 | 1.20732 |
| 200 | 1.4967 |

| **Best-fit values** | |
|---|---|
| $B_{max}$ | 1.86 |
| $K_D$ | 52.02 |
| **Standard Error** | |
| $B_{max}$ | 0.042 |
| $K_D$ | 3.022 |
| **95% Confidence Intervals** | |
| $B_{max}$ | 1.764 to 1.965 |
| $K_D$ | 45.35 to 59.79 |
| **Goodness of Fit** | |
| R square | 0.9984 |

| | EGFR LIR1 |
|---|---|
| GABARAP Y49A L50A [μM] | Response |
| 0 | -0.0043 |
| 0.781 | 0.0673 |
| 1.562 | 0.1469 |
| 3.125 | 0.2756 |
| 6.25 | 0.4638 |
| 12.5 | 0.7549 |
| 25 | 1.1420 |
| 50 | 1.60401 |
| 100 | 2.08511 |
| 200 | 2.53253 |

| **Best-fit values** | |
|---|---|
| $B_{max}$ | 2.96 |
| $K_D$ | 39.03 |
| **Standard Error** | |
| $B_{max}$ | 0.072 |
| $K_D$ | 2.628 |
| **95% Confidence Intervals** | |
| $B_{max}$ | 2.796 to 3.140 |
| $K_D$ | 33.23 to 45.93 |
| **Goodness of Fit** | |
| R square | 0.9976 |

| | EGFR LIR1 |
|---|---|
| GABARAPL1 [μM] | Response |
| 0 | -0.0153 |
| 1.125 | 0.0532 |
| 2.25 | 0.1113 |
| 4.55 | 0.2286 |
| 9.123 | 0.3928 |
| 18.62 | 0.6415 |
| 37.25 | 0.9658 |
| 75 | 1.29569 |
| 150 | 1.65409 |
| 300 | 1.86607 |

| **Best-fit values** | |
|---|---|
| $B_{max}$ | 2.13 |
| $K_D$ | 29.73 |
| **Standard Error** | |
| $B_{max}$ | 0.027 |
| $K_D$ | 1.137 |
| **95% Confidence Intervals** | |
| $B_{max}$ | 2.068 to 2.196 |
| $K_D$ | 27.18 to 32.53 |
| **Goodness of Fit** | |
| R square | 0.9992 |

| | EGFR LIR1 |
|---|---|
| GABARAPL1 Y49A L50A [μM] | Response |
| 0 | -0.0156 |
| 1.125 | 0.1682 |
| 2.25 | 0.3075 |
| 4.55 | 0.5589 |
| 9.123 | 0.8973 |
| 18.62 | 1.3466 |
| 37.25 | 1.8917 |
| 75 | 2.37697 |
| 150 | 2.83305 |
| 300 | 3.19823 |

| **Best-fit values** | |
|---|---|
| $B_{max}$ | 3.406 |
| $K_D$ | 19.07 |
| **Standard Error** | |
| $B_{max}$ | 0.065 |
| $K_D$ | 1.208 |
| **95% Confidence Intervals** | |
| $B_{max}$ | 3.257 to 3.564 |
| $K_D$ | 16.37 to 22.20 |
| **Goodness of Fit** | |
| R square | 0.9974 |

| | PCM1 LIR |
|---|---|
| GABARAP [μM] | Response |
| 0 | 0 |
| 0.195 | 0.195 |
| 0.39 | 0.39 |
| 0.781 | 0.781 |
| 1.562 | 1.562 |
| 3.125 | 3.125 |
| 6.25 | 6.25 |
| 12.5 | 12.5 |
| 25 | 25 |
| 50 | 50 |

| **Best-fit values** | |
|---|---|
| $B_{max}$ | 3.316 |
| $K_D$ | 4.189 |
| **Standard Error** | |
| $B_{max}$ | 0.137 |
| $K_D$ | 0.592 |
| **95% Confidence Intervals** | |
| $B_{max}$ | 3.001 to 3.675 |
| $K_D$ | 2.925 to 5.989 |
| **Goodness of Fit** | |
| R square | 0.9862 |

| | PCM1 LIR |
|---|---|
| GABARAP Y49A L50A [μM] | Response |
| 0 | 0 |
| 0.781 | 0.195 |
| 1.562 | 0.39 |
| 3.125 | 0.781 |
| 6.25 | 1.562 |
| 12.5 | 3.125 |
| 25 | 6.25 |
| 50 | 12.5 |
| 100 | 25 |

| **Best-fit values** | |
|---|---|
| $B_{max}$ | 1.409 |
| $K_D$ | 99.19 |
| **Standard Error** | |
| $B_{max}$ | 0.057 |
| $K_D$ | 6.652 |
| **95% Confidence Intervals** | |
| $B_{max}$ | 1.286 to 1.562 |
| $K_D$ | 84.82 to 117.3 |
| **Goodness of Fit** | |
| R square | 0.9992 |

| | PCM1 LIR |
|---|---|
| GABARAPL1 [μM] | Response |
| 0 | -0.0003 |
| 0.195 | 0.1929 |
| 0.39 | 0.3294 |
| 0.781 | 0.5662 |
| 1.562 | 0.9124 |
| 3.125 | 1.3489 |
| 6.25 | 1.8468 |
| 12.5 | 2.335 |
| 25 | 2.81901 |
| 50 | 3.2468 |

| **Best-fit values** | |
|---|---|
| $B_{max}$ | 3.423 |
| $K_D$ | 4.901 |
| **Standard Error** | |
| $B_{max}$ | 0.09473 |
| $K_D$ | 0.4485 |
| **95% Confidence Intervals** | |
| $B_{max}$ | 3.205 to 3.642 |
| $K_D$ | 3.866 to 5.935 |
| **Goodness of Fit** | |
| R square | 0.9945 |

| | PCM1 LIR |
|---|---|
| GABARAPL1 Y49A L50A [μM] | Response |
| 0 | 0.0009 |
| 0.781 | 0.0150 |
| 1.562 | 0.0259 |
| 3.125 | 0.0383 |
| 6.25 | 0.0627 |
| 12.5 | 0.1145 |
| 25 | 0.2082 |
| 50 | 0.36237 |
| 100 | 0.62008 |
| 200 | 1.00336 |

| **Best-fit values** | |
|---|---|
| $B_{max}$ | 2.363 |
| $K_D$ | 273.3 |
| **Standard Error** | |
| $B_{max}$ | 0.1246 |
| $K_D$ | 21.86 |
| **95% Confidence Intervals** | |
| $B_{max}$ | 2.075 to 2.650 |
| $K_D$ | 222.9 to 323.7 |
| **Goodness of Fit** | |
| R square | 0.9991 |

#### Fig. 4C Bio-Layer Interferometry (BLI) affinity measurements of EGFR LIR1 with GABARAP and Pen8-*ortho*

| | EGFR LIR1 |
|---|---|
| GABARAP [µM] | Response |
| 0 | -0.0081 |
| 1.125 | 0.1081 |
| 2.25 | 0.1894 |
| 4.55 | 0.3376 |
| 9.123 | 0.5630 |
| 18.62 | 0.8780 |
| 37.25 | 1.2865 |
| 75 | 1.74951 |
| 150 | 2.24111 |
| 300 | 2.80478 |

*see Fig. 1B

| Best-fit values | |
|---|---|
| $B_{max}$ | 3.156 |
| $K_D$ | 53.13 |
| **Standard Error** | |
| $B_{max}$ | 0.123 |
| $K_D$ | 6.011 |
| **95% Confidence Intervals** | |
| $B_{max}$ | 2.881 to 3.498 |
| $K_D$ | 40.21 to 70.49 |
| **Goodness of Fit** | |
| R square | 0.993 |

| | | EGFR LIR1 |
|---|---|---|
| GABARAP [µM] | Pen8-ortho [µM] | Response |
| 0 | | -0.0040 |
| 1.125 | 1.7578125 | 0.0415 |
| 2.25 | 3.515625 | 0.0600 |
| 4.55 | 7.03125 | 0.0941 |
| 9.123 | 14.0625 | 0.1372 |
| 18.62 | 28.125 | 0.1810 |
| 37.25 | 56.25 | 0.2565 |
| 75 | 112.5 | 0.35112 |
| 150 | 225 | 0.50093 |
| 300 | 450 | 1.01833 |

*see Fig. 1B

| Best-fit values | |
|---|---|
| $B_{max}$ | 2.482 |
| $K_D$ | 459.9 |
| **Standard Error** | |
| $B_{max}$ | 1.020 |
| $K_D$ | 277.88 |
| **95% Confidence Intervals** | |
| $B_{max}$ | 1.204 to +infinity |
| $K_D$ | 130.2 to +infinity |
| **Goodness of Fit** | |
| R square | 0.9505 |

| | | EGFR LIR1 |
|---|---|---|
| GABARAP [µM] | DMSO [%] | Response |
| 0 | | -0.0198 |
| 0.781 | 0.01 | 0.0363 |
| 1.562 | 0.02 | 0.0591 |
| 3.125 | 0.05 | 0.0842 |
| 6.25 | 0.09 | 0.1147 |
| 12.5 | 0.19 | 0.1572 |
| 25 | 0.38 | 0.2024 |
| 50 | 0.75 | 0.22658 |
| 100 | 1.50 | 0.22482 |
| 200 | 3.00 | 0.19403 |

*DMSO control

| Best-fit values | |
|---|---|
| $B_{max}$ | 2.27 |
| $K_D$ | 45.14 |
| **Standard Error** | |
| $B_{max}$ | 0.07 |
| $K_D$ | 3.723 |
| **95% Confidence Intervals** | |
| $B_{max}$ | 2.111 to 2.450 |
| $K_D$ | 37.04 to 55.21 |
| **Goodness of Fit** | |
| R square | 0.9966 |

#### Fig. S1C Bio-Layer Interferometry (BLI) affinity measurements of EGFR derived phosphopeptides with hATG8 paralogs

| | (pY)EGFR LIR1 |
|---|---|
| GABARAP [µM] | Response |
| 0 | -0.0085 |
| 1.125 | 0.1211 |
| 2.25 | 0.2126 |
| 4.55 | 0.3799 |
| 9.123 | 0.6337 |
| 18.62 | 0.9897 |
| 37.25 | 1.4394 |
| 75 | 1.94058 |
| 150 | 2.43888 |
| 300 | 2.96447 |

| Best-fit values | |
|---|---|
| $B_{max}$ | 3.294 |
| $K_D$ | 46.23 |
| **Standard Error** | |
| $B_{max}$ | 0.107 |
| $K_D$ | 4.447 |
| **95% Confidence Intervals** | |
| $B_{max}$ | 3.051 to 3.570 |
| $K_D$ | 36.55 to 58.60 |
| **Goodness of Fit** | |
| R square | 0.9947 |

| | (pY)EGFR LIR1 |
|---|---|
| GABARAPL1 [µM] | Response |
| 0 | -0.0010 |
| 1.125 | 0.1415 |
| 2.25 | 0.2455 |
| 4.55 | 0.4355 |
| 9.123 | 0.6986 |
| 18.62 | 1.0385 |
| 37.25 | 1.4158 |
| 75 | 1.73477 |
| 150 | 1.9523 |
| 300 | 2.00622 |

| Best-fit values | |
|---|---|
| $B_{max}$ | 2.169 |
| $K_D$ | 19.55 |
| **Standard Error** | |
| $B_{max}$ | 0.019 |
| $K_D$ | 0.614 |
| **95% Confidence Intervals** | |
| $B_{max}$ | 2.126 to 2.213 |
| $K_D$ | 18.18 to 21.02 |
| **Goodness of Fit** | |
| R square | 0.9993 |

| | (pY)EGFR LIR1 |
|---|---|
| GABARAPL2 [µM] | Response |
| 0 | -0.0024 |
| 1.777 | 0.0235 |
| 3.554 | 0.0330 |
| 7.109 | 0.0532 |
| 14.21 | 0.0915 |
| 28.44 | 0.1539 |
| 56.88 | 0.2673 |
| 113.8 | 0.41569 |
| 227.5 | 0.57 |
| 455 | 0.59939 |

| Best-fit values | |
|---|---|
| $B_{max}$ | 0.761 |
| $K_D$ | 98.04 |
| **Standard Error** | |
| $B_{max}$ | 0.031 |
| $K_D$ | 10.768 |
| **95% Confidence Intervals** | |
| $B_{max}$ | 0.6961 to 0.8361 |
| $K_D$ | 76.96 to 125.4 |
| **Goodness of Fit** | |
| R square | 0.9941 |

| | (pY)EGFR LIR1 |
|---|---|
| LC3A [µM] | Response |
| 0 | -0.0288 |
| 2.929 | 0.0641 |
| 5.859 | 0.1407 |
| 11.72 | 0.2689 |
| 23.44 | 0.4294 |
| 46.88 | 0.7288 |
| 93.75 | 1.1318 |
| 187.5 | 1.55749 |
| 375 | 2.00117 |
| 750 | 2.38066 |

| Best-fit values | |
|---|---|
| $B_{max}$ | 2.764 |
| $K_D$ | 135.7 |
| **Standard Error** | |
| $B_{max}$ | 0.05 |
| $K_D$ | 6.979 |
| **95% Confidence Intervals** | |
| $B_{max}$ | 2.650 to 2.887 |
| $K_D$ | 120.1 to 153.3 |
| **Goodness of Fit** | |
| R square | 0.9986 |

| | (pY)EGFR LIR1 |
|---|---|
| LC3B [µM] | Response |
| 0 | -0.0063 |
| 3.906 | 0.0060 |
| 7.812 | 0.0386 |
| 15.63 | 0.1044 |
| 31.25 | 0.1851 |
| 62.5 | 0.3179 |
| 125 | 0.5426 |
| 250 | 0.81411 |
| 500 | 1.16678 |
| 1000 | 1.48066 |

| Best-fit values | |
|---|---|
| $B_{max}$ | 1.968 |
| $K_D$ | 337.9 |
| **Standard Error** | |
| $B_{max}$ | 0.036 |
| $K_D$ | 14.566 |
| **95% Confidence Intervals** | |
| $B_{max}$ | 1.323 to 1.998 |
| $K_D$ | 205.8 to 555.5 |
| **Goodness of Fit** | |
| R square | 0.9992 |

| | (pY)EGFR LIR2 |
|---|---|
| GABARAP [µM] | Response |
| 0 | -0.0077 |
| 1.125 | 0.0099 |
| 2.25 | 0.0243 |
| 4.55 | 0.0487 |
| 9.123 | 0.0918 |
| 18.62 | 0.1681 |
| 37.25 | 0.3016 |
| 75 | 0.5093 |
| 150 | 0.81745 |
| 300 | 1.28761 |

| Best-fit values | |
|---|---|
| $B_{max}$ | 2.56 |
| $K_D$ | 302.3 |
| **Standard Error** | |
| $B_{max}$ | 0.136 |
| $K_D$ | 26.608 |
| **95% Confidence Intervals** | |
| $B_{max}$ | 2.276 to 2.934 |
| $K_D$ | 247.4 to 376.6 |
| **Goodness of Fit** | |
| R square | 0.09985 |

| | (pY)EGFR LIR2 |
|---|---|
| GABARAPL1 [µM] | Response |
| 0 | -0.0081 |
| 1.125 | 0.0392 |
| 2.25 | 0.0663 |
| 4.55 | 0.1297 |
| 9.123 | 0.2172 |
| 18.62 | 0.3459 |
| 37.25 | 0.5028 |
| 75 | 0.67105 |
| 150 | 0.7962 |
| 300 | 0.82962 |

| Best-fit values | |
|---|---|
| $B_{max}$ | 0.9351 |
| $K_D$ | 30.91 |
| **Standard Error** | |
| $B_{max}$ | 0.013 |
| $K_D$ | 1.373 |
| **95% Confidence Intervals** | |
| $B_{max}$ | 0.9065 to 0.9649 |
| $K_D$ | 27.92 to 34.22 |
| **Goodness of Fit** | |
| R square | 0.9988 |

| | (pY)EGFR LIR2 |
|---|---|
| GABARAPL2 [µM] | Response |
| 0 | -0.0088 |
| 1.777 | 0.0053 |
| 3.554 | 0.0053 |
| 7.109 | 0.0152 |
| 14.21 | 0.0245 |
| 28.44 | 0.0430 |
| 56.88 | 0.0806 |
| 113.8 | 0.14257 |
| 227.5 | 0.2096 |
| 455 | 0.26338 |

| Best-fit values | |
|---|---|
| $B_{max}$ | 0.3834 |
| $K_D$ | 199.2 |
| **Standard Error** | |
| $B_{max}$ | 0.014 |
| $K_D$ | 15.831 |
| **95% Confidence Intervals** | |
| $B_{max}$ | 0.3540 to 0.4181 |
| $K_D$ | 167.1 to 239.2 |
| **Goodness of Fit** | |
| R square | 0.9978 |

| | (pY)EGFR LIR2 |
|---|---|
| LC3A [µM] | Response |
| 0 | -0.0158 |
| 2.929 | 0.0129 |
| 5.859 | 0.0442 |
| 11.72 | 0.0950 |
| 23.44 | 0.1468 |
| 46.88 | 0.2648 |
| 93.75 | 0.4413 |
| 187.5 | 0.6588 |
| 375 | 0.90798 |
| 750 | 1.16002 |

| Best-fit values | |
|---|---|
| $B_{max}$ | 1.495 |
| $K_D$ | 229.7 |
| **Standard Error** | |
| $B_{max}$ | 0.033 |
| $K_D$ | 12.298 |
| **95% Confidence Intervals** | |
| $B_{max}$ | 1.421 to 1.577 |
| $K_D$ | 202.6 to 261.0 |
| **Goodness of Fit** | |
| R square | 0.9988 |

| | (pY)EGFR LIR2 |
|---|---|
| LC3B [µM] | Response |
| 0 | -0.0169 |
| 3.906 | -0.0117 |
| 7.812 | 0.0064 |
| 15.63 | 0.0335 |
| 31.25 | 0.0511 |
| 62.5 | 0.0987 |
| 125 | 0.1785 |
| 250 | 0.29326 |
| 500 | 0.44105 |
| 1000 | 0.59362 |

| Best-fit values | |
|---|---|
| $B_{max}$ | 0.8956 |
| $K_D$ | 511.2 |
| **Standard Error** | |
| $B_{max}$ | 0.033 |
| $K_D$ | 39.09 |
| **95% Confidence Intervals** | |
| $B_{max}$ | 0.5487 to 0.9476 |
| $K_D$ | 259.9 to 854.3 |
| **Goodness of Fit** | |
| R square | 0.9982 |
